# High Throughput Viability Assay for Microbiology

**DOI:** 10.1101/2023.01.04.522767

**Authors:** Christian T. Meyer, Grace K. Lynch, Dana F. Stamo, Eugene J. Miller, Anushree Chatterjee, Joel M. Kralj

## Abstract

Counting viable cells is a universal practice in microbiology. The colony forming unit (CFU) assay has remained the gold standard to measure viability across disciplines; however, it is time-intensive and resource-consuming. Herein, we describe the Geometric Viability Assay (GVA) that replicates CFU measurements over 6-orders of magnitude while reducing over 10-fold the time and consumables. GVA computes a sample’s viable cell count based on the distribution of embedded colonies growing inside a pipette tip. GVA is compatible with gram-positive and -negative planktonic bacteria, biofilms, and yeast. Laborious CFU experiments such as checkerboard assays, treatment time-courses, and drug screens against slow-growing cells are simplified by GVA. We therefore screened a drug library against exponential and stationary phase *E. coli* leading to the discovery of the ROS-mediated, bactericidal mechanism of diphenyliodonium. The ease and low cost of GVA evinces it can accelerate existing viability assays and enable measurements at previously impractical scales.

## Main

The colony forming unit (CFU) assay is the gold standard for enumerating viable cells in microbiology labs across the world [1–6]. The CFU assay combines simplicity with readily available reagents to achieve an enormous dynamic range, commonly measuring between 1 and 100,000,000 viable cells in a sample. Viability measurements are critical in numerous contexts spanning food safety [7], functional genomics [8–10], and drug combination discovery against persister cells [2, 11]. However, measuring viability across numerous conditions using the CFU assay is time- and resource-intensive while generating a significant amount of plastic waste [4, 12].

Prior approaches to increase the scale of viability measurements included 1) increasing the speed with robotic liquid handling and imaging [1, 4, 13]; 2) decreasing the amount of pipetting by using viability stains [14] or microfluidics [15]; or 3) using cell growth to estimate the initial number of viable cells post-treatment [3]. The most commercially successful alternative to the CFU assay is the Spiral Plater method [16] which deposits the sample in an Archimedes spiral on a solid medium plate. However, none of these approaches combines the simplicity, low cost, dynamic range, and versatility as simply diluting cells and then growing them on solid media.

Herein, we developed a new viability assay, called the Geometric Viability Assay (GVA). GVA calculates the CFUs in a sample based on the axial position of embedded colonies that form in a cone. Intuitively, the probability of a colony forming at the tip of the cone is less than near the base. Analytically, we find this probability is proportional to the squared perpendicular distance of the colony to the cone tip. By measuring the position of a few colonies in the cone and utilizing the derived probability function, the total number of colonies in the entire cone can be computed with high precision. By leveraging the latent information encoded in the colony distribution, GVA accurately quantified the number of viable cells in a sample ranging from 1 cell to 1,000,000. This dynamic range was accomplished using a cone universal in microbiology—the pipette tip. In summary, GVA 1) measures viability over *>*6 orders of magnitude; 2) does not depend on the cell’s growth or lag phase; 3) minimizes consumables; and, 4) reduces operator time by over 30-fold compared to the drop CFU assay. Combined this enabled throughputs of up to 1,200 viability measurements per researcher per day.

## Results

### The Geometric Viability Assay

The most time- and resource-intensive step of the classic drop CFU is the dilution series that must be run to count individual colonies across several orders of magnitude. We reasoned the geometry of a cone could create a dilution series in a single step as the cross section at the tip is less than the cross section near the base. Analytically, the probability of a colony forming at any point along the cone’s axis proportional to the cross-sectional area at that point (Fig. 1a, cyan circle). This probability is defined as the probability density function (PDF) equal to

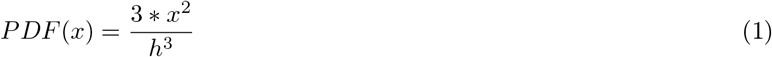

where *x* is the perpendicular distance from the tip along the x-axis and *h* is the total length of the cone (Figs. 1a, S1a-c; see Supplemental Materials for derivation). Equation (1) is applicable for arbitrary cones or pyramids which are axially symmetric (Fig. S1d). The total CFU concentration in the cone can be estimated by

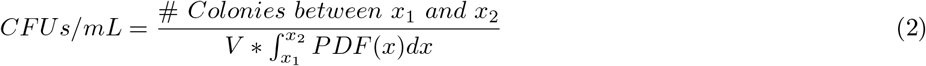

where (*x*_1_,*x*_2_) are the positions of the first and last colony in the counted sub-volume and *V* is the volume of the cone. Thus, the highest CFU density resolvable is proportional to the dynamic range of the PDF. In contrast to a cylinder or a wedge, the cone achieves the maximum dynamic range in the PDF by changing shape in all 3 dimensions (Fig. 1b). Importantly, this probability does not depend on the radial (*y, z*) position of a colony within the cone, only on the perpendicular distance from the tip along cone’s axis (*x*).

**Figure 1:**
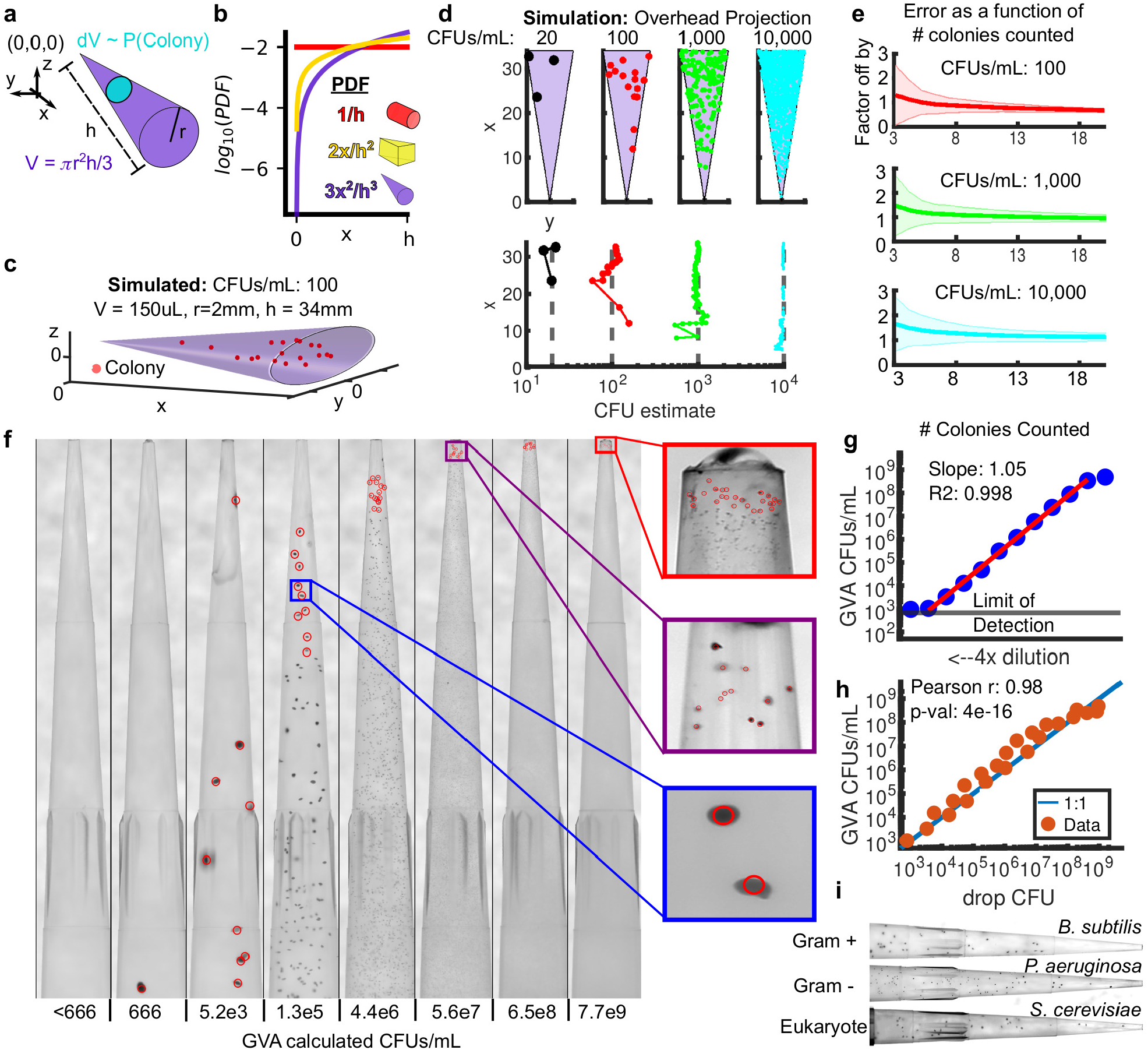
The Geometric Viability Assay (GVA). a) The probability of a colony forming at a distance *x* from the tip of the cone is proportional to the infinitesimal volume *dV* (cyan circle) divided by the total volume *V* (purple cone). Analytically, this ratio is the Probability Density Function (PDF) as a function of *x* (see Supplemental Materials for derivation). b) The PDF for a cylinder (red), wedge (yellow), and cone (purple) as a function of the axial distance (x). c) Simulation of the colony distribution in a cone. d) Estimating the total CFUs/mL based on the position of colonies in the cone. (top) Shown are the distributions of colonies for 4 simulations spanning 20 to 10,000 CFUs/mL density. The volume of each cone is the same as in panel c. (bottom) GVA estimate of the CFUs/mL as a function of the included colonies and their x positions. e) The factor the GVA calculation differs from the correct value as a function of the number of colonies in Equation (1). Shaded errorbars represent 1 standard deviation in 1000 simulations. Colors match simulations in panel d. f) Dilution series of *E. coli* embedded in 150 *µ*L 0.5% LB-agarose in p200 pipette tips. Red circles correspond to colonies counted using a custom semi-automated segmentation software. g) *E. coli* CFUs/mL calculated using GVA for a 4x dilution series. Points are the mean of 4 replicates. Mean calculated after taking the log. Red line is the linear regression fit to dilution series. A slope of 1 on a log-log plot is expected if the GVA estimate scales linearly with dilution. h) The drop CFU and GVA estimates are significantly correlated over 6 orders of magnitude. i) GVA performed on gram-positive, gram-negative, and eukaryotic cells (see Fig. S4a for quantification)

We simulated colony distributions in a cone for different CFUs/mL (Fig. 1c,d). As expected, the more CFUs in the cone, the more colonies are found near the tip (Fig. 1d, top panel). The CFUs/mL estimate quickly converges to the correct value (gray dotted line) as more colonies’ position are included in Equation (2), regardless of the colony density (Fig. 1d, bottom panel). Remarkably, the CFU estimate is off by less than a factor of 2 from the correct value in 97% of simulations based only on the positions of the first 10 colonies, even if there are over 10,000 colonies in the cone (Fig. 1e, S1f). This rapid convergence to the correct value is the same regardless of the CFU concentration. Therefore, by leveraging the information encoded in the geometry of the cone, it is not necessary to count all the colonies to accurately calculate the colony density. This concept is analogous to a 3D hemocytometer; by counting a subset of colonies within a defined volume, the total concentration can be computed using probabilities.

To test the theory, we used a cone ubiquitous in microbiology—the pipette tip. The first experiment was a dilution series using stationary phase *Escherichia coli* (BW25113). CFUs/mL of stationary phase *E. coli* are known to be approximately 10^9^ CFUs/mL after overnight growth [17]. Cells were serially diluted and then each dilution was treated as a sample of unknown concentration of viable cells. Each “sample” was fully mixed with melted LB agarose (cooled to ≤ 50°C) to a final agarose concentration of 0.5%. Triphenyltetrazolium chloride (TTC) was included in the melted agarose to increase the colony contrast. The agarose was allowed to solidify in the tip before the tip was ejected into an empty tip rack (See Methods). The agarose-containing pipette tips were then incubated overnight at 37°C and imaged the following day using a custom build optical setup with a mirrorless Canon camera (Fig. 1f, see Fig. S2 for optical configuration, Supplemental Movie 1 for GVA protocol overview). In agreement with our simulations, the distribution of colonies that form in the tip was predictable based on the PDF across > 6 orders of magnitude (Fig. 1g, slope ∼ 1). Remarkably, the final colony size decreased with increasing cell density which prevented colony overlap even at high densities. Comparing the same batch of cells using GVA and the traditional drop CFU assay showed the two approaches are significantly correlated (Fig. 1h, Pearson r=0.98, p-val=4e-16, see Fig. S3 for example drop CFU plate).

GVA was used to count other gram-negative (*Pseudomonas aeruginosa, Salmonella typhimuirium, Pseudomonas putida*) and a gram-positive bacterial strain (*Bacillus subtilis*) as well as eukaryotic yeast cells (*Saccharomyces cerevisiae*) (Figs. 1i, S4a). Enclosing the colonies in a pipette tip facilitated handling pathogenic strains because a bleach wash could kill all contaminating cells on the outside of the tip without affecting colony growth inside the tip (Fig. S4b). Viability in *E. coli* biofilms over time was also tested with GVA (Fig. S4c,d). Finally, we tested the potential of GVA for rapid quantitation of non-model bacterial species. Human-associated biome viability measurements were conducted using GVA (Fig. S5). Vigorously swabbing 24 locations (Fig. S5a) revealed a large dynamic range of microbial concentrations capable of growth in LB (Fig. S5b). Growing sample replicates at different temperatures revealed temperature-selective growth for different biomes (Fig. S5c). These experiments necessarily underestimate the number of bacteria in these biomes as many human commensals are unculturable. However, because GVA uses solid growth media, the same selective culturing techniques developed over the last 100 years for standard petri dish plating can be leveraged in GVA while also enabling high throughput surveillance of culturable biomes.

For all cultures, samples were embedded in 0.5% agarose melted in culture medium: LB for the bacteria and YEPD for the yeast. Embedded bacterial colonies also grew in other media such as Meuller Hinton Broth or M9 minimal medium. Using 3D printed molds to embed the colonies in a square pyramid (Fig. S6a,b), we confirmed the GVA approach was applicable to geometries other than cones, as predicted (Fig. S6c-j).

We next investigated how the dynamic range and accuracy of GVA depended on the optical configuration using a low cost camera system—an iPhone with a commercial macro lens. We designed a pipette tip holder that positioned a single tip in front of an iPhone rear camera and a macro lens (Fig. 2a,S7). Calibration revealed the pixel size of the iPhone was 13.7 microns compared to 6.6 microns for the Canon EOS camera with 100mm f/2.8 macro lens (Fig. S2). We reasoned the smaller pixel size and lower electron depth in the iPhone camera would reduce the smallest possible colony detected as compared to the mirrorless camera. As expected, comparing images taken with the Canon camera with the iPhone demonstrated colonies at the highest CFU concentrations were no longer resolvable on the iPhone (Fig. 2b). Comparing the GVA-calculated CFUs/mL for the same pipette tips of an *E. coli* dilution series using both the iPhone and the Canon camera, we measured a reduction in dynamic range of 64X on the iPhone as compared to the Canon camera (Fig. 2c). However, GVA remained highly linear for nearly 5 orders of magnitude (green line, slope=1.04, R^2^=0.99) with the iPhone configuration. The correlation between the CFU counts for iPhone and Canon configurations on the same pipette tips was 0.99 (Fig. 2d). Therefore, we found GVA is accurate regardless of the optical configuration, but the dynamic range is set by the maximum camera resolution.

**Figure 2:**
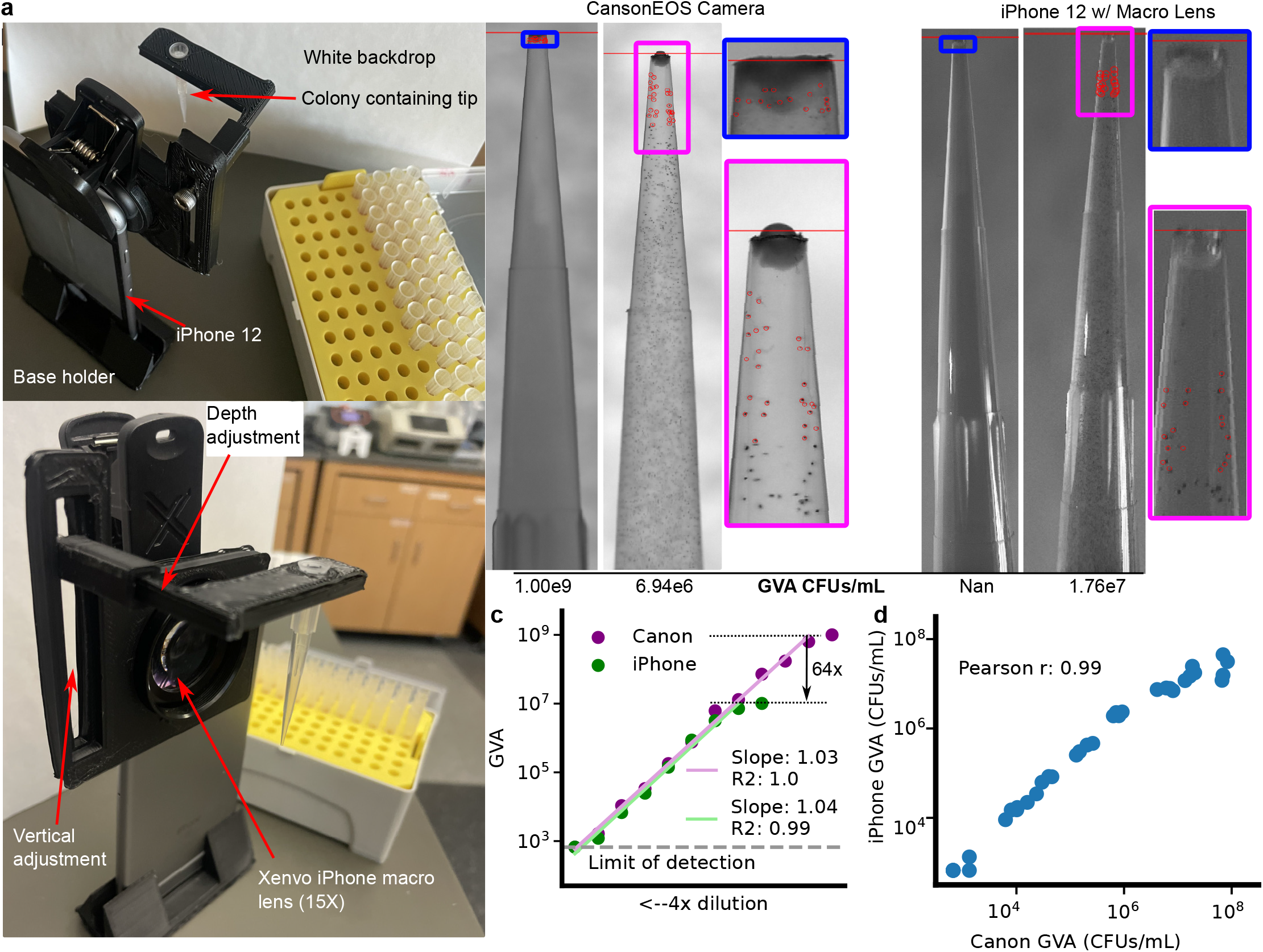
GVA dynamic range, but not accuracy, depends on the optical configuration. a) Picture of assembled pipette tip holder on an iPhone 12 with a Xenvo macro lens. The pipette images are taken in front of a white backdrop (paper) with ambient illumination. b) Example images of the same 2 pipette tips using the Canon EOS with 100 mm f2.8 macro lens (left) or the iPhone 12 with Xenvo macro lens (right). The GVA calculated CFUs/mL are reported below. Selected colonies for GVA calculation are circled. c) Dynamic range of the iPhone GVA. *E. coli* were diluted 4X and embedded in pipette tips. After incubation, the same tips were imaged with the iPhone camera with macro lens (green) and the mirrorless camera (purple). Points are the mean of 4 replicates calculated after taking the log. Green and purple lines are the linear regression fit to the dilution series. d) Pearson correlation between iPhone GVA and professional camera for all pipettes where colonies could be counted using both. Correlation coefficient calculated in log-space.

The main advantage of GVA is the more than 10x reduction in time, reagent cost, and plastic waste as compared to the drop CFU or Spiral Plater methods (Fig. 3). The Spiral Plater is the most common commercial alternative for the CFU assay utilizing a specialized instrument to dilute the sample along an Archimedes spiral [16]. In order to measure the time savings of GVA, we compared 3 steps of viability assays including the preparation of solid growth media (Fig. 3b), diluting/plating 96 conditions (Fig. 3c), and imaging/counting of the colonies (Fig. 3d). The largest time savings was in the plating step. The drop CFU took 3 hours to manually plate 96 conditions. Current Spiral Plater instruments are reported to take 30 seconds per plate, corresponding to 96 conditions in 48 minutes. GVA took 5 minutes corresponding to a 36X savings in time for plating. GVA was also faster in the time for preparation than both the Spiral Plater and drop CFU approaches. The time for imaging and counting the colonies was the fastest on the Spiral Plater according to the manufacturer-reported time using an automated colony counter. GVA semi-automated colony counting took a similar amount of time to manual colony counting for the drop CFU when including the time for image acquisition, pipette tip segmentation, and user-guided colony detection. In total, using the current instrumentation, a single researcher measured the viability of 1,200 conditions in a day.

**Figure 3:**
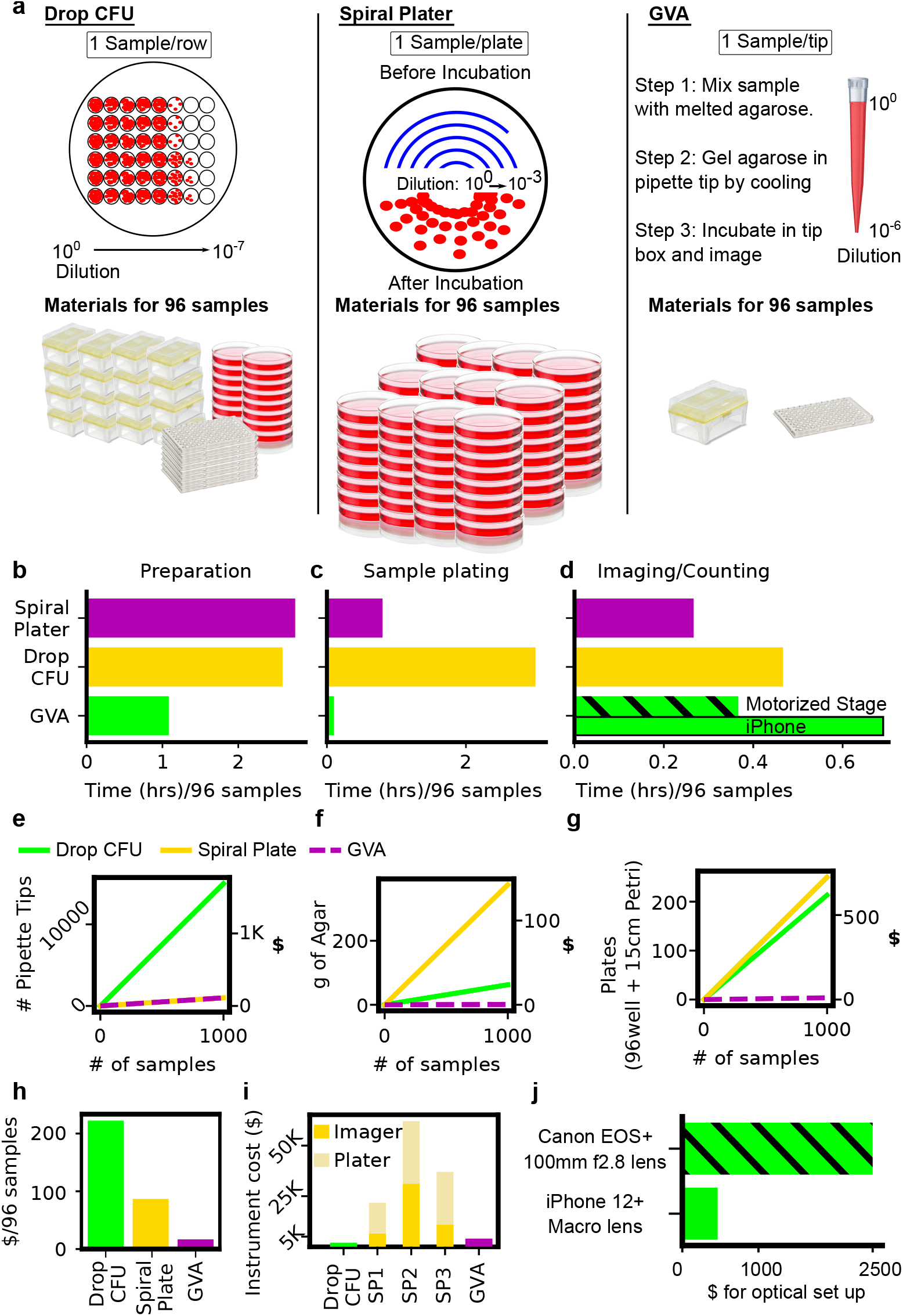
GVA reduces the time and materials of viability measurements by over 10-fold. a) (left) Schematic of a drop CFU assay and required materials for 96 samples assuming tips are changed for each dilution step. (middle) A Spiral Plater spreads a sample in an Archimedes spiral on a solid media plate. The spiral results in decreasing sample volume as a function of radial distance with a reported 3-log dynamic range. One petri dish is required per sample. (right) GVA uses a single pipette tip to run a 6-order dilution series. b-d) Time comparisons for different techniques. b) Time required to prepare solid growth media. The preparation time for the Spiral Plater and drop CFU includes: 1) autoclaving the agar; 2) cooling post autoclave; 3) plate pouring; and 4) an plate cooling. GVA melts agarose in a microwave which is subsequently equilibrated in a warm bath for 1 hour prior to starting. c) Sample plating from a 96-well plate. Time for the Spiral Plater assay sample plating based on industry-reported value. Drop CFU was timed by an expert user using a 12-channel pipette and changing tips at every dilution and plating step. d) Time required for quantification of 96 samples. Spiral Plater time is based on industry-reported value using an automated colony counter. GVA time includes imaging (7 min for Canon with motorized stage and 30 min for iPhone), image preprocessing and tip segmentation (5min), and semi-automated colony counting (10min) for 96 pipette tips. The drop CFU colonies were counted and recorded manually. e) Number and cost of pipette tips as a function of sample count for the three different techniques. See Supplemental Table 1 for cost estimates. f) Amount of agar required as a function of sample count. 25mL of 1.5% agar per 15cm petri dish was assumed for the drop CFU and Spiral Plater assays. 200 *µ*L of 0.5% agarose per tip was assumed for the GVA. g) Number of 96-well and petri dishes per condition. h) Estimated total cost in consumables per 96 samples of the three methods. GVA cost is $0.17/sample. i) Instrument costs. Based on quotes for a Spiral Plater (SP) and automated imaging system from 3 manufacturers. GVA instrument cost included the Canon camera and 100mm f/2.8 macro lens. j) The difference in instrumentation cost for the Canon and iPhone optical configurations.

We next compared the reagent savings and plastic waste reduction of the three approaches. In the drop CFU assay, since each sample must be diluted and then separately transferred to an agarose pad, 15 pipette tips per sample is standard for our laboratory protocol (Fig. 3e)[18]. In GVA, a single pipette tip is used per sample amounting to a 15x savings in pipette tips over the drop CFU (Fig. 3e). In the Spiral Plater assay, a petri dish with solid growth medium is required per condition (Fig. 3g). Compared to the Spiral Plater method, the plastic required is reduced from a petri dish to a pipette tip. Summing the cost of pipette tips, agar, and culture plates at the time of writing, we found the drop CFU was the most expensive in consumables costing an average of $222 per 96 samples compared to the Spiral Plater and GVA which cost $87 and $17, respectively (Fig. 3h, see Supplemental Table 1 for pricing rationale). The savings in consumables of the Spiral Plater is offset by the substantial instrument costs (Fig. 3i, see Supplemental Table 1). Costs were calculated from quotes for 3 Spiral Platers and automated imaging systems solicited from three distributors. The instrument costs for both the GVA and the drop CFU included an electronic, multichannel pipette. Additional instrumentation costs for the GVA depended on the optical configuration (Fig. 3j) which were at least an order of magnitude less than the Spiral Plater systems.

In summary, our analysis showed GVA substantially reduced operator time, instrument and reagent costs, and the carbon footprint of viability assays.

We next investigated the robustness of GVA. We first measured the count noise between 4 technical replicates across CFU concentrations ranging between 10^2^ and 10^7^ CFUs/mL (Fig. 4a,b). Noise was calculated using the coefficient of variation (COV) between replicates. Across all measured CFU concentrations, the GVA noise is less than or equal to the noise of the drop CFU assay for both the Canon and iPhone optical configurations. As with the drop CFU assay, the GVA noise is heteroskedastic, increasing as the number of colonies decreases as expected for a Poisson process.

**Figure 4:**
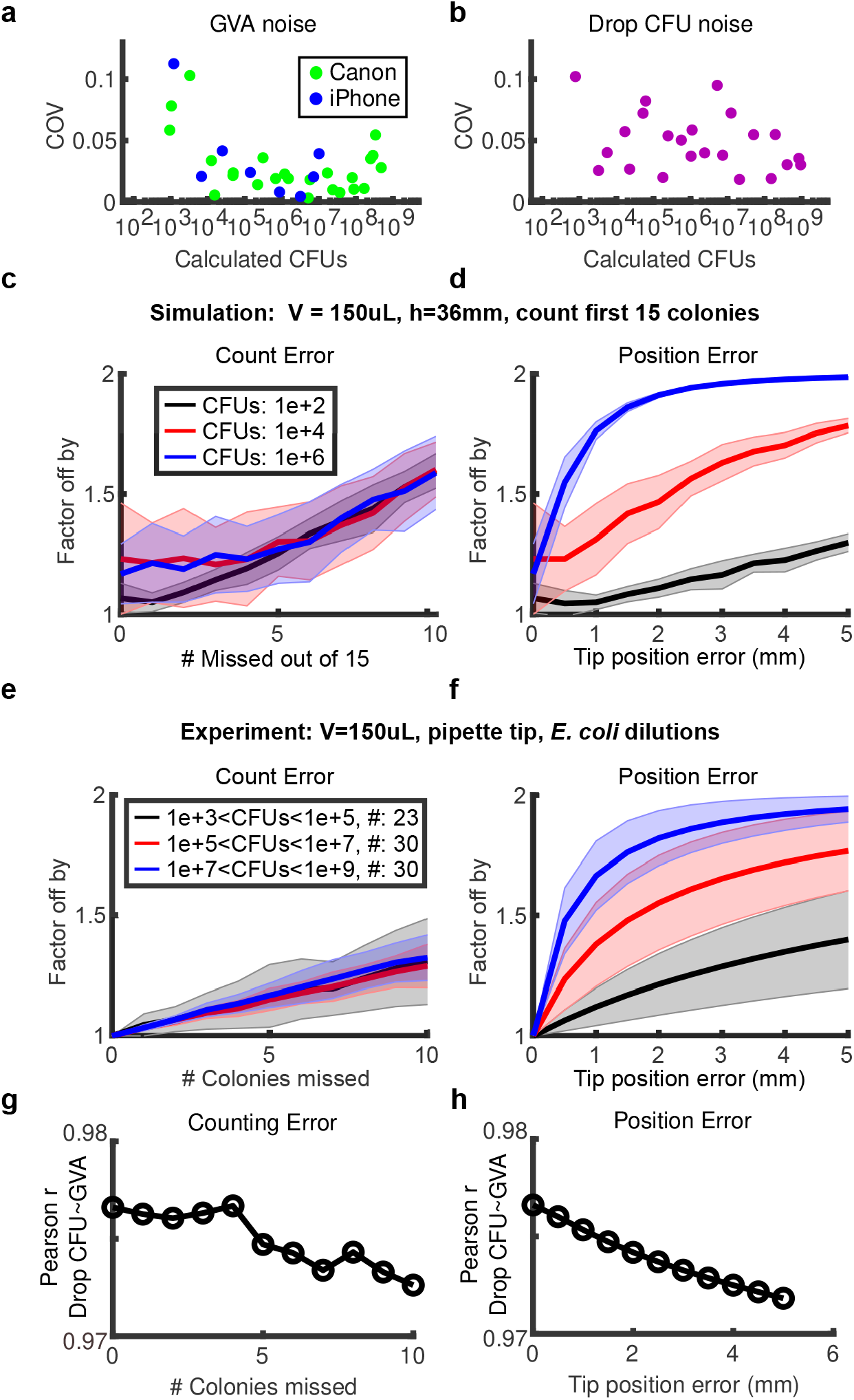
GVA has a low noise profile and is robust to missing colonies or tip position errors. a,b) Coefficient of Variation (COV) between 4 technical replicates for different number of CFU concentrations for GVA using the Canon or iPhone optical configuration (a) and drop CFU (b). c,d) The factor the GVA calculation differs from the correct value as a function of the number of missed colonies (c) or error in tip position (d) in simulated results (see Methods). Shaded error bar is the standard deviation in 1000 simulations. e,f) Same error calculations for experimental data. Error bars represent the standard deviation between all the pipette tips (#) included in each bin. g,h) Correlation between the GVA and the drop CFU assay as a function of counting and position errors.

After confirming GVA’s low technical noise, we investigated the impacts of two types of real-wold errors on GVA calculations: missing colonies and uncertainty in the position of the cone tip. These errors were examined using both simulated and experimental data. Predictably, as the number of missed colonies increases, the error increases (Fig. 4c,e) though the fractional error is the same in all seeding densities. Remarkably, eliminating 10 out of 15 counted colonies in the simulated data resulted in estimates within a factor of 2, regardless of the initial CFU concentration. This robustness was recapitulated in the experimental data and is in agreement with the observation that the position of only 5 colonies is sufficient to calculate the CFUs/mL within a factor of 2 on average (Fig. 1e). For pipette tip position errors, the GVA calculations at high CFU concentrations are more sensitive to misidentification of the tip position than low cell concentrations (Fig. 4d,f blue versus black lines). Nevertheless, missing the tip position by 10% (4 mm for a 36 mm cone) still resulted in an estimate within a factor of 2 from the correct value in both simulations and experiments. Finally, the correlation between the drop CFU and the GVA (Fig. 1h) decreased modestly from 0.98 to 0.97 for combinations of missing up to 10 colonies and missing the tip position by 4 mm (Figs. 4g,h, S8). These simulated and experimental data highlight the robustness of GVA.

In total, our analyses find GVA is accurate and robust, retaining sensitivity over comparable ranges to the gold standard drop CFU while reducing the cost and time.

### High throughput viability screening against stationary phase *E. coli*

Previous studies have found slow growth is a non-inheritable form of antibiotic tolerance buying time for viable cells to develop genetic resistance [19]. Slow-growing cells commonly have reduced metabolic activity [2] and DNA replication [20] as compared to exponentially growing cells. As a result, slow-growing cells are refractory to antibiotics targeting DNA synthesis (fluoroquinolones) [21], protein translation (aminoglycosides) [22], and cell wall biogenesis (beta-lactams) [23]. Growth-dependent tolerance can only be observed by measuring viability but the tedium and cost of the drop CFU assay limits extensive profiling. Using GVA, we directly compared the viability of exponentially growing cells and stationary phase cells to different doses of three antibiotics for varying amounts of time. In total, we tested 3 antibiotics at 6 different concentrations for 5 different durations against stationary and exponential cells, in duplicate, for a total of 360 viability measurements (Fig. 5a,b). This data was acquired by a single researcher in one day using only 4 tip boxes. Stationary phase cells were more resistant to ciprofloxacin, carbenicillin, and gentamicin. Particularly, for carbenicillin, there was less than a 10-fold decline in viability of stationary cells treated with 100 *µ*g/mL carbenicillin for 24 hours, as compared to a 10,000 fold decrease in exponential cells. Treating exponential cells with 10 *µ*g/mL carbenicillin showed no change in the number of colonies during the first 6 hours, followed by an increase in viable cells after 24 hours treatment indicative of a slowly-expanding, drug-tolerant pool (Fig. 5b). Ciprofloxacin at 10 *µ*g/mL had a biphasic pharmacodynamic profile with initial bactericidal activity within an hour resulting in a 10-fold reduction in viability for both stationary and exponentially growing cultures. However, this activity stabilized through 6 hours and a second phase of killing was achieved by 24 hours. Gentamicin at 10 *µ*g/mL required a full 24 hours to achieve more than a 10-fold reduction in stationary phase cell viability. For untreated cultures, we observed the concentration of exponentially growing cells increased till a peak concentration of *∼* 10^9^ CFUs/mL at 6 hours (Fig. S9). Once in stationary phase, the number of viable cells declined over time as previously reported [17]. These data exemplified the utility of GVA for measuring the efficacy of treatments agnostic to growth rate.

**Figure 5:**
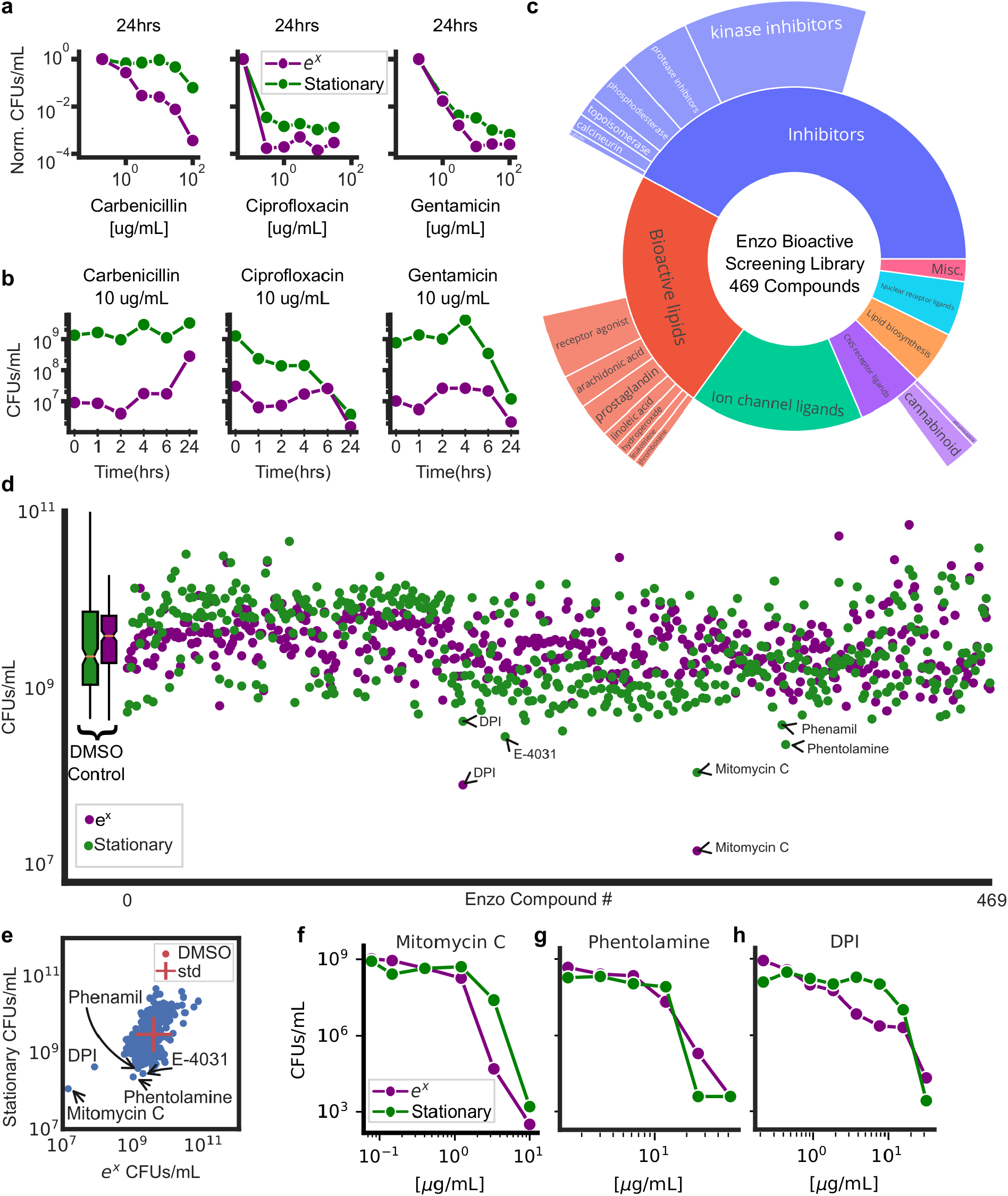
GVA screening of the Enzo library identifies DPI as active against stationary phase *E. coli*. a) Dose-response of 3 antibiotics for stationary and exponential (*e*^*x*^) cultures after 24 hours of treatment. Each point is the mean of duplicate measurements. CFUs/mL were normalized to an untreated control. b) Viability over time for stationary and exponential cells at one concentration of antibiotic. c) Drug classes of the Enzo Bioactive Screening Library. Size of donut wedge is proportional to drug class representation. Targets of each class and relative representation depicted on the outer ring. d) Absolute viability of stationary (green) and exponentially growing (purple) cells after 24 hours of treatment with Enzo library. Each condition was run in duplicate and the mean taken in log-space. e) Scatter plot of stationary phase versus exponential phase from the screen. The standard deviation of DMSO controls are depicted with a red cross. Selected hits are annotated. f,g,h) Mitomycin C (DNA crosslinker), phentolamine (*α*-adrenergic antagonist), and DPI (NADPH oxidase inhibitor) dose responses in stationary and exponential cultures.

To explore the GVA technique’s potential for high throughput viability measurements, we screened the ICCB Enzo Bioactive library (469 compounds) against stationary and exponentially growing cultures (Fig. 5c,d). The Enzo library has a wide breadth of chemical matter including bioactive lipids, small molecule inhibitors, and ion channel ligands (Fig. 5c) and spans the structural diversity of larger libraries like the Maybridge HitFinder library of approximately 14,000 compounds (Fig. S10a). Viability of BW25113 *E. coli* treated with the Enzo library was measured in both exponential and stationary phase. Including controls and removing pipette errors, 2267 conditions were measured. The equivalent screen using the drop CFU or Spiral Plater assays would have required 355 tip boxes or 2267 petri dishes, respectively. GVA required 24 tip boxes. No edge effects were observed for either stationary or exponential plates (Mann-Whitney U test, p-val>0.05, Fig. S10b). Average differences among drug classes were modest (Fig. S10c, p-val > 0.001 ANOVA, p-val corrected for multiple hypothesis testing) and none significantly different than the control (p-val*>*0.01, Pairwise Tukey Test). Five compounds were selected for follow up verification (mitomycin C, phentolamine, E-4031, phenamil, and diphenyliodonium) corresponding to a ∼ 1% hit rate. Mitomycin C is a known antibiotic acting through DNA cross-linking. As expected, we found it is more active against exponentially growing cells compared to cells in stationary phase (Fig. 5f). Phentolamine is an *α*-adrenergic receptor antagonist. Phentolamine has previously been shown to block norepinephrine- and epinephrine-induced growth in *E. coli* putatively by antagonizing *α*-adrenergic-like receptors [24]. We found at high concentrations (20 *µ*g/mL) stationary cells were more sensitive to the effects of phentolamine than exponentially growing cells (Fig. 5g) corroborating the differential sensitivity observed in the screen. E-4031 and phenamil did not have any dose-dependent effect on viability (Fig. S11). Finally, we found diphenyleneiodonium (DPI), a promiscuous NADPH Oxidase (NOX) inhibitor [25], to be active against both stationary and growing cultures (Fig. 5h). Previous studies have identified DPI as possessing antimicrobial characteristics [26, 27]; however, the mechanism of DPI bactericidal activity remains unknown. We were intrigued by DPI’s bacteriocidal activity as it reduces Reactive Oxygen Species (ROS) in eukaryotes by inhibiting NOXs [25] which is in contrast to the mechanism of many antibiotics which increase ROS pools [28–30].

In order to investigate the bactericidal mechanism of DPI, we first examined *E. coli* ROS levels upon treatment with DPI. ROS levels were determined with the fluorescent CellROX dye which measures cytoplasmic superoxide [31]. Single cell fluorescence was measured over time after treatment with a lethal DPI dose and compared to an untreated control (Fig. 6a). As expected, DPI substantially decreased ROS reaching the nadir around 75 minutes after the drug was added (Fig. 6a compare blue and yellow lines, Supplemental Movie 2). The depth and duration of the ROS reduction was proportional to the DPI concentration (Fig. S12a). Surprisingly, this decrease was followed by a rapid spike in ROS. In contrast to DPI, ciprofloxacin treatment resulted in monotonically increasing levels of ROS (Fig. 6a, orange line). Increased levels of ROS underlie ciprofloxacin’s bactericidal activity [32]; therefore, we next investigated if the ROS spike induced by DPI also underlies its bactericidal activity. We compared DPI sensitivity of stationary phase cells in aerobic versus anaerobic environments. DPI was less active in anaerobic cultures (Fig. 6b), similar to gentamicin or ciprofloxacin (Fig. S12b,c). This data suggested high levels of ROS are part of the bactericidal mechanism of DPI, despite it initially decreasing ROS. In further support of this, adding a ROS scavenger also reduced DPI efficacy (Fig. S12d).

**Figure 6:**
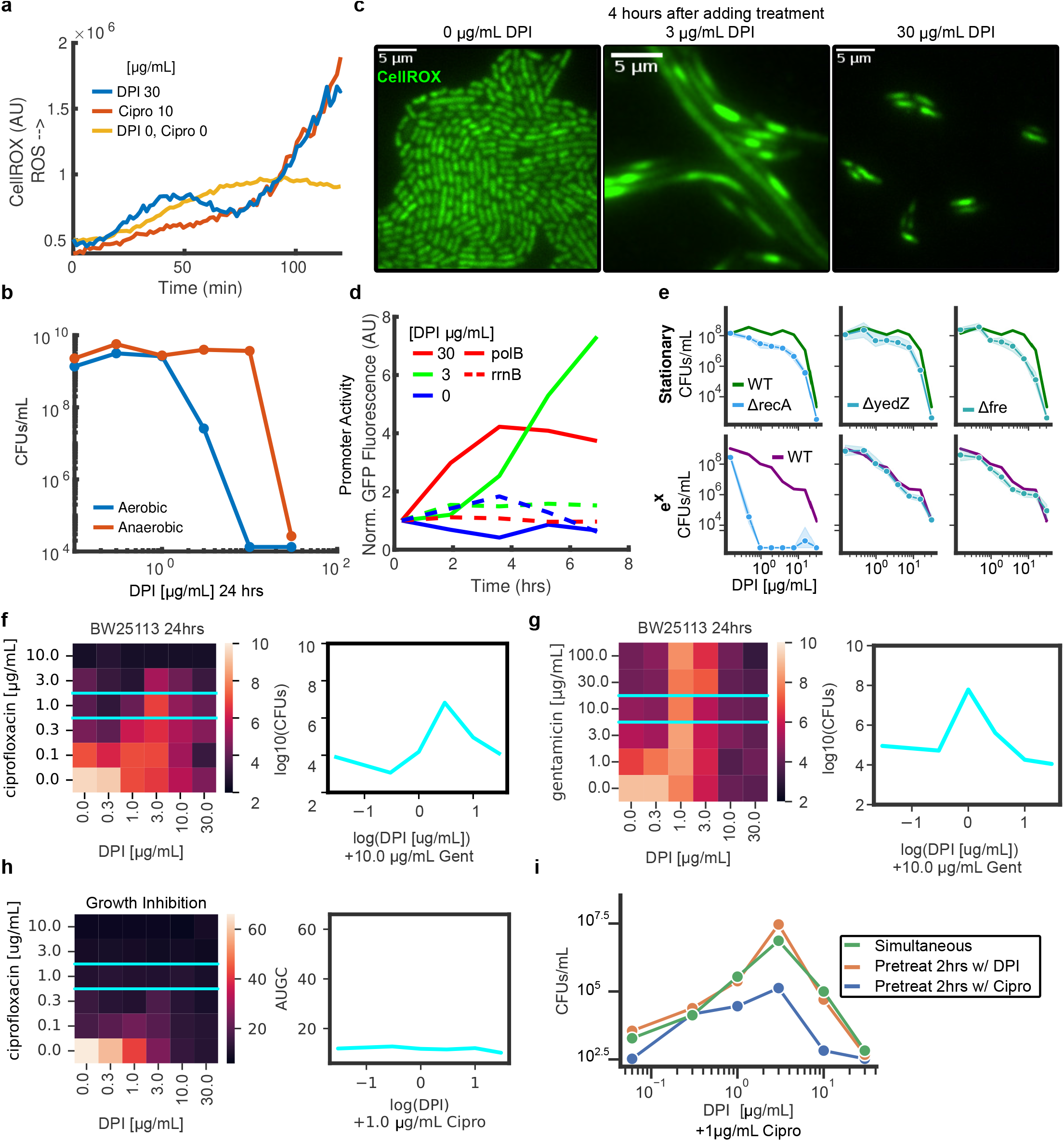
DPI generates ROS, activates the SOS response, and antagonizes ciprofloxacin. a) Median, single-cell CellROX signal as a function of time for DPI (blue), ciprofloxacin (orange), and an untreated control (yellow). b) Efficacy of DPI in aerobic and anaerobic conditions. See Fig. S12b,c for ciprofloxacin and gentamicin. c) Images of live *E. coli* cells stained with the CellROX dye for three DPI concentrations 4 hours after adding DPI. Brightness and contrast is the same for all images. See Supplemental Movie 2. d) Measurement of *polB* and *rrnB* promoter activity normalized to t=0. e) DPI dose response for *E. coli* knockout mutants treated during stationary (top panels) or exponential growth (bottom panels). The dose response for the wild-type (WT) cells is depicted in green or purple, respectively. Shaded errorbars equal to the standard deviation in logspace between 3 replicates. See Fig. S14 for other mutants. f) GVA checkerboard assay for DPI combined with ciprofloxacin at 24 hours. Each square in the heatmap was the mean of duplicate conditions. Colorbar correspond to the log10(CFUs/mL) for each dose combination. Left panel shows the dose response for DPI plus 1 *µ*g/mL ciprofloxacin (cyan). See Fig. S15 for full time series. g) GVA checkerboard assay for DPI combined with gentamicin at 24 hours. h) Growth inhibition checkerboard for DPI and ciprofloxacin. Optical density was measured for each condition over 8 hours and the integrated area under the growth curve (AUGC) is depicted (colorbar). i) Dose response curves for temporally staggered combinations. All treatments lasted for 24 hours total. Pretreated conditions were treated for 2 hours with a single drug followed by 22 hours with both drugs.

Intermediate DPI concentrations altered ROS levels but maintained viability when measured with GVA. We examined the cell morphology after 4 hours of treatment with less than 10 *µ*g/mL DPI and observed the formation of bacterial filaments (Fig. 6c, Supplemental Movie 2). Filamentation is a classic hallmark of SOS activation [33] and increases in ROS are an established SOS activator [34]. Therefore, we wondered if DPI was activating SOS. We examined the promoter activity of genes downstream of lexA using the PEC GFP-promoter library [35]. LexA is a master transcriptional repressor of genes in the SOS regulon such as *polB, dinB, dinG*, and *yjiH*, and is auto-catalytically degraded by activated recA. We observed persistent, dose-dependent induction of the *polB* promoter compared to a ribosomal protein control (*rrnB*) (Fig. 6d, solid versus dashed lines). The highest promoter activity corresponded to an intermediate dose of DPI (3 *µ*g/mL) where filamentation was observed. We also observed a DPI-dependent increase in *dinB, dinG*, and *yjiH* promoter activity (Fig. S13). The *lexA* promoter, which is self-repressed, also increased activity within 90 minutes of DPI addition.

Because SOS activity reduces the efficacy of other bactericidal agents [36], we predicted that recA-mediated SOS activation was critical for maintaining viability in the presence of DPI. As predicted, *recA* knockouts were more susceptible to DPI in both stationary and exponential phases of growth (Fig. 6e), though the increased DPI potency was more pronounced in exponentially growing cells. In contrast, knocking out other DNA repair enzymes, redox repair enzymes, or ROS scavengers did not substantially change the potency of DPI in either growth phase (Fig. S14). Knocking out *yedZ* and *fre*, genes recently identified as part of a NOX-like system in bacteria [37], modestly increased the potency of DPI against stationary cells indicating these proteins are unlikely to be the main target of DPI in *E. coli* (Fig. 6e). Therefore, our data showed DPI activated SOS, and that SOS activation enhanced cell viability.

We therefore wondered if DPI would antagonize other antibiotics whose efficacy is reduced by the SOS response. Such antagonism has been observed in combinations of ciprofloxacin with metronidazole, a redox-active prodrug known to activate SOS [38]. To test for antagonism, we measured viability in a time-resolved, checkerboard assay using GVA (Figs. 6f, S15). In the checkerboard assay, DPI was combined with either ciprofloxacin or gentamicin across a 6 × 6 dose matrix. The ease of GVA enabled sampling the checkerboard over time, resulting in a complete pharmacokinetic profile of the drug-drug interaction. DPI antagonized both ciprofloxacin and gentamicin against stationary phase *E. coli* increasing the viability 1,000-fold as compared to either drug alone after 24 hours treatment (Fig. 6f,g). This antagonism was not observed in a growth inhibition assay (Fig. 6h) emphasizing the value of viability data when investigating drug-drug interactions. DPI antagonism of ciprofloxacin and gentamicin was also observed in *S. typhimuirium* (Fig. S16). Cells pretreated with DPI for 2 hours before adding ciprofloxacin further increased protection, while pretreating with ciprofloxacin for 2 hours reduced DPI’s antagonistic effects (Fig. 6i).

In total, we found DPI initially decreased ROS followed by a ROS burst which enhanced its bactericidal effects. As expected with previous studies of ROS lethality [34], the potency of DPI depended on SOS-activation mediated via recA. By activating SOS, DPI led to an increase in drug tolerance to fluoroquinolones and aminoglycosides as revealed by temporal viability checkerboards.

## Discussion

One of the most surprising features of GVA was how well the theory enabled accurate viability estimates in practice regardless of the optical configuration. In simulations and experiments, errors in the colony count and tip position did not substantially alter CFU estimations when considering the experimental dynamic range. Furthermore, pipette tips are not perfect cones; small imperfections in manufacturing were clearly visible at high magnifications. Despite these real world variances—using an imperfect cone, selecting a few colonies, and approximating the tip location—GVA still reproducibly and accurately calculated CFU concentrations across 6 orders of magnitude. This robustness emerges from utilizing the latent information encoded in a colony’s position.

Another unexpected feature of GVA was the observation of self limiting colony size depending on the CFU density. As the concentration of colonies increased, the commensurate decrease in colony size preserved colony discreteness even for dense samples. Colony size, in the strains tested, plateaued after overnight incubation and did not change over several additional days. The physiological basis of this phenomenon remains unknown, though we speculate it could be due to quorum sensing, nutrient limitation, or mechanically-inhibited growth. However, the self-limiting colony growth in 3D may not be universally true of microbes, which would limit the applicability of GVA.

GVA suffers from the same culturability limitations as the drop CFU [39, 40]. Additionally, it is unknown how many organisms that grow in 2D will not grow in 3D or vise versa; however, GVA worked for all commonly used laboratory strains tested as well as more complex samples such as biofilms and human-associated biomes samples. Because GVA uses the same growth substrate as historic 2D culture techniques (e.g. solid media), we anticipate many of the tricks that have evolved to selectively culture different strains in petri dishes will be transferable to GVA. Growth in 3D may also alter the antibiotic sensitivity due to mechanosensitive changes in physiology [41, 42]; therefore, rigorous testing is required to compare MIC values for 2D versus 3D plating. Finally, the transient thermal shock of the current protocol using agarose did not impact viability of the tested strains, but could be a non-trivial perturbation for certain species. In such cases, the use of other hydrogels which crosslink via chemical reaction (*e*.*g*. sodium alginate) may be appropriate.

In both the drop CFU and Spiral Plater methods, the incubation time remains a rate limiting step, commonly taking at least overnight for visible colonies to emerge. For GVA, incubation is also a rate limiting step; however, we achieved colony detection across all CFU concentrations within 8 hours for *E. coli*. This improvement in time to detection is due to the unique optical configuration, the presence of a staining dye, and the 3D geometry which maximizes light scattering. Decreasing time further could be achieved with the use of fluorescent imaging. Though we expect time to detection for *E. coli* to be the experimental floor, this proof-of-concept data suggest GVA could be a means to reduce the time of clinical antibiotic sensitivity profiling.

In total, we find the GVA approach to substantially reduce the time and reagents required for measuring cell viability compared to the established drop CFU assay while maintaining the same dynamic range, quantitative nature, and versatility across different species that has made the drop CFU assay the gold standard for viability measurements in microbiology.

## Methods

### Strains and growth conditions

*E. coli* strain BW25113 was used unless otherwise noted in the text. This strain was acquired from the Yale Coli Genetic Stock Center. *E. coli* was grown in LB (Sigma Aldrich) at 37°C in a shaking incubator. *B. subtilis* strain W168 was a kind gift from the Garner lab and was grown in LB at 37°C in a shaking incubator. *P. putida* strain KT2440 was a kind gift from Jacob Fenster and was grown in LB at 30°C in a shaking incubator. *S. typhimurium* strain SL1344 was a kind gift from the Corrie Detweiler and was grown in LB at 37°C in a shaking incubator. *S. cerevisiae* strain BY4741 was a kind gift from Roy Parker and was grown in YEPD at 30°C in a shaking incubator. *P. aeruginosa* strain PA01 was a kind gift from the Zemer Gitai and was grown in LB at 37°C in a shaking incubator. Knockouts were selected from the Keio collection (Dharmacon). The PEC promoter library in *E. coli* was acquired from Dharmacon (PEC3877).

All bacterial and yeast strains were streaked onto an agar plate with appropriate antibiotic selection if required (kanamycin for Keio and PEC strains). These plates were kept for 1 month in a 4°C refrigerator. Individual colonies were then selected and grown overnight in 3-5 mL cultures in 12 mL culture tubes with appropriate antibiotic selection if required. Each colony selected was considered a biological replicate. Multiple measurements of the same culture were considered technical replicates.

### Antibiotic treatments

Antibiotic treatments were typically performed in 96 well plates with a 12 channel electronic pipette. For stationary phase treatments, bacterial cells were grown overnight (≥ 16 hours) in a shaking incubator (180RPM). For *P. putida* only, cells were grown for 2 days. Upon entering the stationary phase, cells were distributed into a 96 well flat-bottom plates with 100 *µ*L of cells per well. Drug treatments at 1000x were plated into a separate 96 well-round bottom plate. A 100 nL pin transfer was used to dilute the drug plate into the cell plate at a 1:1000 ratio. This plate was then placed into a shaking incubator for the experimental time.

To measure antibiotic treatments in the exponential phase, overnight culture was diluted 1:1000 into fresh LB. This culture was then placed into the incubator for 2 hours. After this incubation, the cells were then distributed to the 96 well plate followed by drug treatment.

### Drop CFU assay

Drop CFU assays were performed similar to the method described in [18]. Briefly, in a 96 well plate, 90 *µ*L was added to all wells except row A. Into row A, a 100 *µ*L volume of sample solution was added. From row A, 10 *µ*L of cells was taken and added into row B, followed by 3 mixes. This process was repeated from B to C, until the final dilution on row H corresponding to a 1e-7 dilution from the original sample. Pipette tips were changed for each row to reduce sample carry over. From each column of the dilution series, 3 *µ*L drops were transferred onto an LB-agar pad. Once all the liquid was absorbed into the agar (typically 15-30 minutes), the agar plates were inverted and placed into a 37°C standing incubator overnight. Counting the next morning was performed by hand. The first dilution with individually resolvable colonies was used to count and multiplied by the corresponding dilution factor.

### Embedding for GVA

The goal for embedding was to have a uniformly mixed sample in liquid hydrogel that would quickly solidify the 3D mold. We used 0.5% agarose as a convenient hydrogel that would solidify quickly and prevent cell motility once solidified. Pipette tips (200 *µ*L, VWR universal) were most commonly used as a reproducible and cheap 3D geometry scaffold.

1. *Preparing the agarose solution*. A 0.66% agarose solution was prepared in the cell medium of choice. We found the color of LB and YEPD did not affect the imaging in the pipette tips. Agarose (0.66 g) was added to a 100 mL volume of LB and microwaved until completely dissolved. A careful watch was maintained during the heating to ensure it did not boil over. Upon full dissolution, the liquid was placed in a 50°C heat bath to maintain in liquid state until ready to use. At this stage, tetrazolium chloride (TTC, 25 *µ*g/mL final concentration) was added to the LB-agarose from a 1000x stock for all bacteria experiments. Respiring bacteria reduce tetrazolium to water-insoluable formazan which stains the colonies red.
2. *Preparing the cells*. A fresh 96 round-bottom plate was prepared by adding 50 *µ*L of LB or YEPD to each well. The sample plate with the cells and drugs was removed from the shaking incubator, and a pin transfer tool (2 *µ*L hanging drop, VP409) was used to transfer 2 *µ*L of the treated cells into the 50 *µ*L LB plate. If conducting a time-course experiment, the sample plate was then placed back into the shaking incubator.
3. *Embedding*. To embed, we found an electronic multichannel pipettor was the most convenient for high numbers of samples. We typically used a 12 channel P200 (Eppendorf explorer, 4861000724). The following items were gathered before pouring the liquid agarose into a reservoir: the 96-well plate with 20 *µ*L samples (from step 2), a box of autoclaved P200 pipette tips, an empty P200 tip box filled with ice water, an empty P200 tip box with 2 mL water in the bottom to hold the embedded cells. At this point, the liquid agarose was poured into a 100 mL reservoir for easy use with the multichannel pipette. Using the pipet and mix function on the pipetor, 150 *µ*L of the LB agarose solution was taken from the reservoir, and mixed twice with 1 row of the sample plate (200 *µ*L final volume, 0.5% final agarose concentration, 1:100 dilution from the sample plate). After mixing, 150 *µ*L was taken into the same pipette tips avoiding bubble formation. These tips were then placed into the ice bath for 6 seconds to ensure the hydrogel was solidified to plug the tip. Then the tips were ejected into the empty pipette tip box. This process was repeated for all 7 additional rows in plate. Using 150 *µ*L and the 1:100 dilution from the original sample gave a lower limit of 667 CFUs/mL.
4. *Incubation*. Upon completion of the embedding process, the tip box with the LB-agarose-cell suspension is left at room temperature for *∼*30 minutes to ensure the agarose is fully solidified. The tips were then moved into a standing incubator overnight for the colonies to grow. We found that the colonies did not change size after overnight incubation so that cells could be imaged up to 4 days post embedding as long as they were maintained in a hydrated environment.

### Drug screens

A screen was performed with the ICCB Enzo Bioactive hits library (Enzo, BML-2840-0100). An overnight culture of 60 mL LB was grown to stationary phase with *E. coli*. The next morning, 60 *µ*L of the overnight culture (stationary phase) was added to a fresh 60 mL of LB and grown for the 2 hours in the shaking incubator (exponential phase). The cells were then dispensed into 100 *µ*L volumes into 96 well plates.

### Biofilm Growth and Treatment

MG1655 *E. coli* strains were used for biofilms. Overnight cultures were diluted 1:10^5^ in LB. Biofilms were seeded in a U-bottom 96 well plate and grown for 48 hours at 37°C in a stationary incubator. For temporal experiments, a separate plate was used for each timepoint and biofilms were dispersed at the indicated times. Reported time represent the number of hours after the initial 48 hour incubation. To disperse the biofilms, non-adhered cells were aspirated, wells were washed with PBS, and fresh PBS was added to the wells. The plate was covered with foil plate seals (VWR, 60941-126) and put on a plate shaker at 3000 rpm for 30 minutes. Dispersed cells were diluted 10^4^ and GVA was performed. A crystal violet stain was used to confirm proper dispersal; any replicates that were not fully dispersed were discarded.

### Imaging GVA tips

Imaging took place on a custom instrument (Fig. S2) or an iPhone 12 (Fig. 2). For the custom instrument, a mirrorless commercial camera (Canon EOS RP) with a 1:1 macro lens (Canon, f/2.8 100 mm) was used to obtain high quality images that could resolve the smallest colonies. For the iPhone, the parts were designed in FreeCAD and then 3D printed with PLA using a Lulzbot Taz Pro FDM printer. Model files are available via GitHub. All pieces could fit on the print bed in a single print. Print bed adhesion was increased using a glue stick before printing. The print bed temperature was set to 70°C for all layers and the nozzle temperature was set to 225°C. Print speed was set to 10 mm/sec for initial layers and then increased to 30 mm/sec for subsequent layers. Post printing, the depth channel (green in Fig. 2a), was tapped with a 8-32 bit. After the holder was assembled on the Xenvo macro lens with the wide field lens removed, the tip was positioned in front of a white backdrop and imaged with ambient illumination using the iPhone’s autofocus function. 3 images per tip were taken and the tip most in focus was selected before processing using the Matlab app.

The digital camera was mounted above a light box that provided even illumination. The light box was then moved by a stepper stage so that 12 tips could be imaged automatically (3 tips per field of view, 4 fields of view). The light box consisted of a Styrofoam box that was covered by a transparent acrylic sheet (McMaster Carr, #8560K257). A white paper was attached to the underside of the acrylic to act as a diffuser. The inside of the Styrofoam box was lined with foil (Reynolds). A high intensity cyan LED (Luxeon Rebel, 3Up) was placed on a heatsink inside the box and was powered with a constant current driver (BuckBlock, 2100 mA). The Styrofoam light box was mounted onto a stepper motor stage (Thorlabs, LTS300). The camera was mounted using the tripod’s 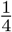 ”-20 screw threads onto a z-translator (Thorlabs, MT1) which was affixed to a right angle plate (Thorlabs, AP90). The Z-positioner was used to set a distance such that 3 pipette tips could be imaged in one field of view, and the macro lens was used to bring them into focus. With our camera, this corresponded to a pixel size of 5.8 *µ*m (Fig. S2c). To place the tips onto the light box, a broken 12 channel P200 head was used. This made loading and unloading samples easy using the spring release while also providing a standard orientation for the tips.

Images were collected with a custom Labview script to control the camera and the stepper stage. Labview called a separate program, digiCamControl (www.digicamcontrol.com) to access camera functions and acquire images. Typical camera settings used a shutter speed of 1/100 s, aperture 6.3, and ISO 100. At each field of view, 5 images were collected followed by a stage movement to the next 3 pipet tips (27 mm). The images were stored directly on the instrument computer as high resolution .jpg files. Using this instrument, a typical experiment of 96 tips could be imaged in *∼*7 minutes.

### Image processing

The goal of the image processing was to identify and extract individual pipet tips from the collected images and identify individual colonies. These were broken into two steps which were performed sequentially. Matlab (Mathworks, R2021b) was used for all image processing analyses. The developed app can be used without a Matlab license using a compiled version specific to the user’s operating system.

#### 1) Pipette tip segmentation

All images from a given field of view were converted to a 16-bit grayscale image. The green channel from the images were summed and that image was used for downstream analyses. The overall orientation of the image was calculated to ensure that each tip was oriented perpendicular to the x-axis. Due to small variations in the tip loading onto the light box, this was necessary to accurately calculate the colony distance from the pipette tip. The Hessian (fibermetric.m) of the image was calculated and convoluted with a horizontal line to locate the angle of the tips. The image was then rotated (imrotate.m) by this angle to orient the pipettes vertically in the image. To identify the x-pixels corresponding to the pipette tip, the Hessian was again calculated from the rotated image. From the middle of the image, a convolution of a single line at different angles was used to calculate the left and right boundaries of the pipette tip. These lines were then extended to the bottom of the pipette tip to locate the left and right boundaries of the tip. Each of the three wells was then saved into a cell array.

#### 2) Semi-automated segmentation

Colonies were segmented using a semi-automated, custom script in Matlab. From the extracted image of the pipette tip, the user selected one of 4 different segmentation routines corresponding to the varying sizes of colonies in the pipette tip. The first routine segmented the entire pipette tip while the last segmentation algorithm zoomed into 1/7th of the full tip and segmented the first 30 colonies. Segmentation was done using Matlab’s Image Processing Toolbox. Subsequently, the user could curate the automated segmentation adding missed colonies or removing erroneous colonies.

The colony count and position of the first and last colonies was used in eqs. (1) and (2) to calculate the GVA estimate of the CFUs/mL. For the error analysis, the factor the GVA estimate differed from the correct value was calculated according to: Factor off by = 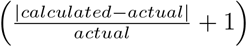. This approach to error calculation takes into account the large dynamic range of possible CFUs/mL.

### Microscopy measurements

For all microscopy experiments, cells from overnight cultures were diluted 1:100 in minimal media (PMM) and shaken for 2 hours at 37°C to ensure cells had exited lag phase. After 2 hours of growth, 2 *µ*L of dilute cell culture was added to the top of a cooled, 200 *µ*L 2% low melt agarose pad with CellROX dye (5*µ*M). The agarose pad was molded to fit in 96-well square bottom plates (Brooks Automation, MGB096-1-2-LG-L). After 10 minutes of drying, the pad with affixed cells was inverted and pressed into the bottom of an imaging plate. Fields of view (FOV) were selected manually on the microscope. After FOVs were selected and before the imaging started, the drug was added on top as done previously [18, 43]. We have previously found the drug diffuses through the pad on the order of minutes [43].

Imaging took place using a Nikon Ti2 inverted microscope running the Nikon Elements software package. Fluorescent excitation was achieved with a laser source (488 nm and 561 nm) using a high-angle illumination to minimize the out-of-focus background. All images were acquired with a 40x, NA 0.95 air objective. Images were acquired on an sCMOS camera (Hamamatsu, ORCA-Fusion) camera.

Image processing was done in Matlab (Mathworks, R2020a) and followed the general scheme described in [18]. Briefly, the illumination profile for all images was estimated from the average of 50 images per FOV. Morphological opening and blurring were used to broaden the illumination pattern before correcting the images. After illumination correction, the jitter in the movie was removed by aligning each sequential frame using a fast 2D Fourier transform implemented in Matlab. The background was locally subtracted based on an estimation of the background computed using morphological image opening before segmentation.

Segmenting cells was done using the Hessian-based *fibermetric* routine implemented in Matlab which is specific for identifying tubular structures. Segmented regions were included only if they met a minimum area and intensity threshold which were manually selected based on the camera and laser settings. To remove rare segmented debris, the mean Euclidean distance of each cell from all other cells in a multi-dimensional feature space was calculated and objects which were in the 95th percentile or above in average distance were removed [43]. A cell’s position in the feature space was defined by its segmented area, perimeter, major/minor axis lengths, and circularity extracted using Matlab’s *regionprops* command.

## Acknowledgments

The authors thank Ian Peck and Paul Koenig of the Bioserve Space Technologies machine shop for their useful input on design of the chips. The authors thank Zachary Berriman-Rozen for helpful early discussions. The authors thank Corrie Detweiler for helpful discussions regarding GVA’s application to screening and Calvin Ewing for his input on optimizing GVA for use in different labs. Finally, the authors thank Aaron Whiteley who inadvertently sparked the idea for using pipette tips.

## Author Contributions

conceptualization, C.T.M. and J.M.K.

methodology, C.T.M., G.K.L, D.F.S., and J.M.K.

formal analysis, C.T.M., E.J.M., and J.M.K.

investigation, C.T.M., G.K.L, D.F.S., E.J.M, and J.M.K.

data curation, C.T.M., E.J.M., and J.M.K.

writing—original draft preparation, C.T.M.

writing—review and editing, C.T.M., G.K.L, E.J.M., A.C., and J.M.K.

software, C.T.M., E.J.M., and J.M.K.

visualization, C.T.M., D.F.S., and J.M.K.

supervision and funding acquisition, A.C and J.M.K.

all authors have read and agreed to the published version of the manuscript.

## Data and Code Availability

The code and data required for recreating the figures will be made available via GitHub pending publication.

## Supplemental Materials

## 1 Supplemental Movies

Supplemental Movie 1: Protocol for GVA sample embedding. weblink

Supplemental Movie 2: Live cell imaging of CellROX stained cells imaged under agarose pad with different concentrations of DPI. Pad is made with PMM. Time (HH:MM) annotated in the upper left. DPI was added on top of pad at start of video. weblink

## 2 Supplemental Tables

Supplemental Table 1: Pricing for viability measurement consumables and Spiral Plater instrumentation. weblink

## 3 Supplemental Figures

**Figure S1:**
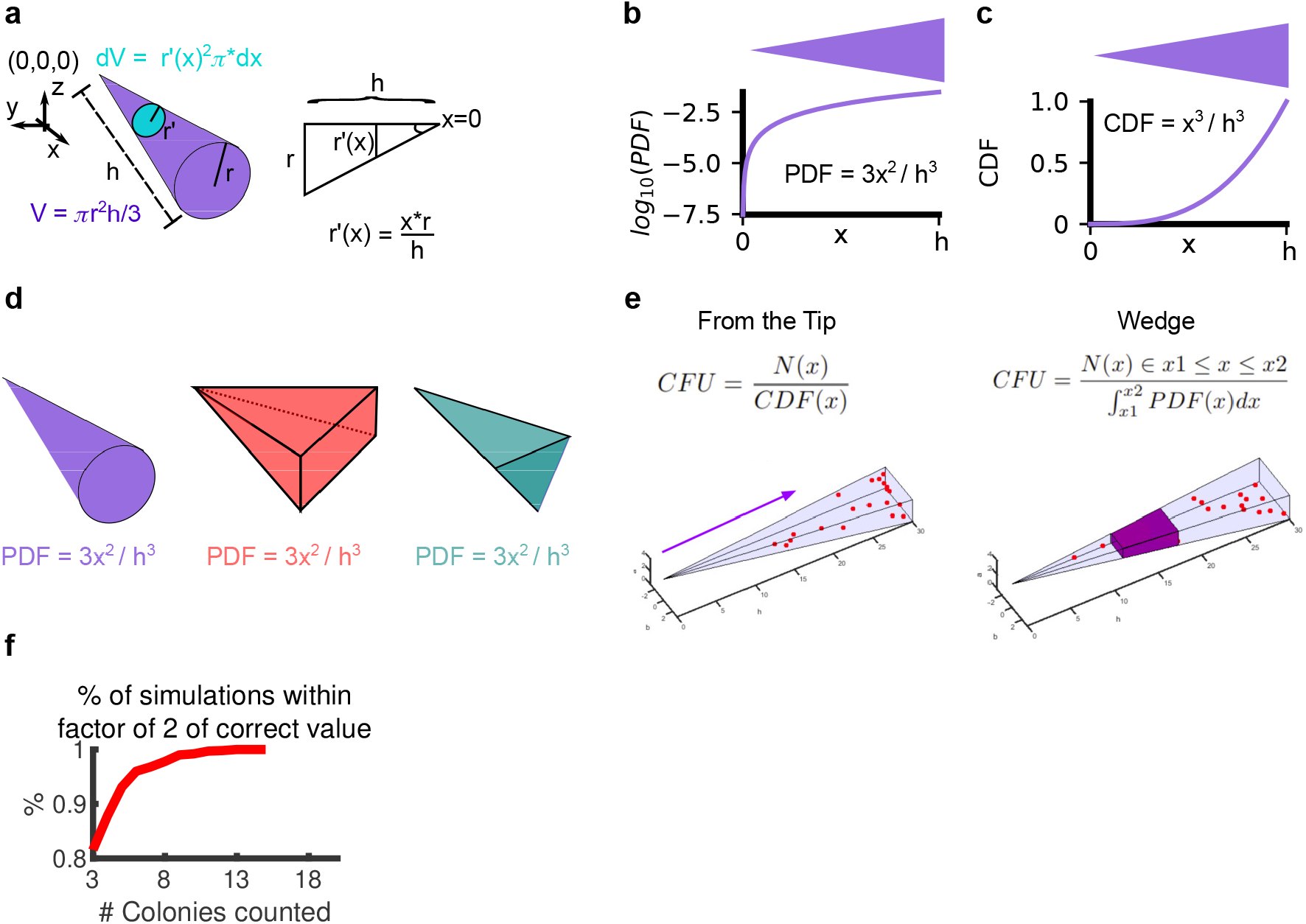
Derivation of a cone’s PDF. a) The volume of the infinitesimal *dV* divided by the total volume *V* corresponds to the probability of finding a colony as a function of *x*. The radius of the infinitesimal (*r*^*′*^(*x*)) is a function of the radius of the cone’s base (*r*) divided by the height of the cone (*h*) times *x* according to trigonometry. b) The PDF of the cone as a function of x. Overhead projection of cone is depicted above. c) The cumulative density function (CDF) as a function of x. d) The PDF is the same for axially symmetric cones such as a square (red) and triangle (turquoise) pyramids. e) Two equivalent ways of calculating the number of CFUs in the wedge using either the CDF (left) or PDF (right). *N* (*x*) is the number of colonies counted. f) Percentage of simulations with the GVA calculated CFUs/mL within a factor of 2 of the correct value as a function of the number of colonies used for the GVA calculation. 1000 simulations used to calculate percentage. See Figure 1c for simulation parameters.

**Figure S2:**
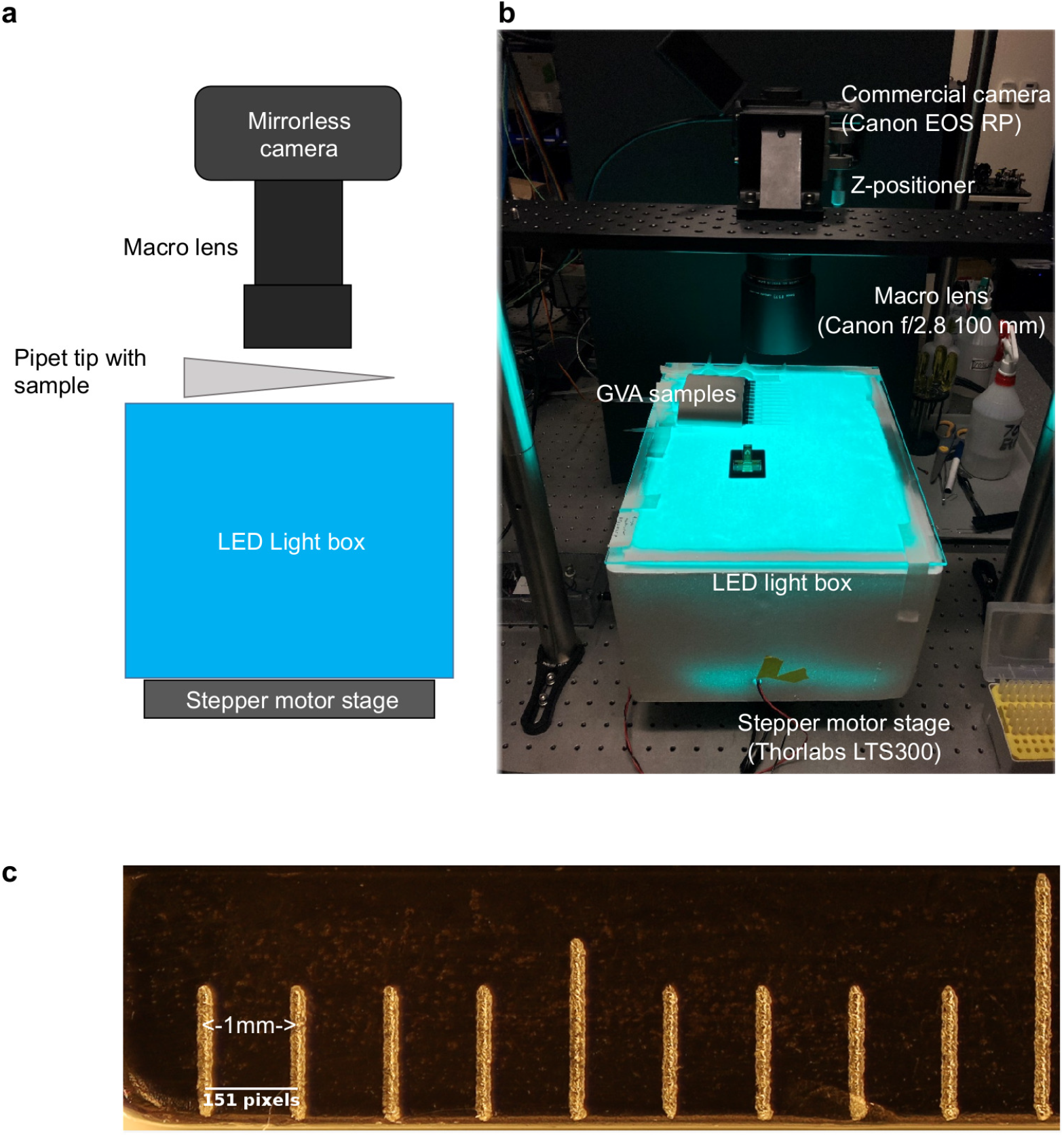
Optical configuration. a) Schematic of optical configuration for imaging the pipette tips containing agarose. A mirrorless camera with a macro lens is positioned above the tips at the focal plane. Addition of a z-positioner stage helps in fine tuning the focus. Pipettes are illuminated transversely using an LED light box with a diffuser. This box is mounted on a stepper motor to allow for imaging 12 pipettes at a time. The stepper motor and the camera are simultaneously controlled via LabView software. b) Picture of optical configuration. A cyan light was used to maximize the contrast of the TTC counterstain. A styrofoam box functions as a reflective light box and a sheet of paper as a diffuser. The GVA samples are positioned using a 12 channel pipetter and imaged using a Canon EOS RP camera with a f/2.8 100mm macro lens. c) Pixel resolution for this configuration is 6.7 microns.

**Figure S3:**
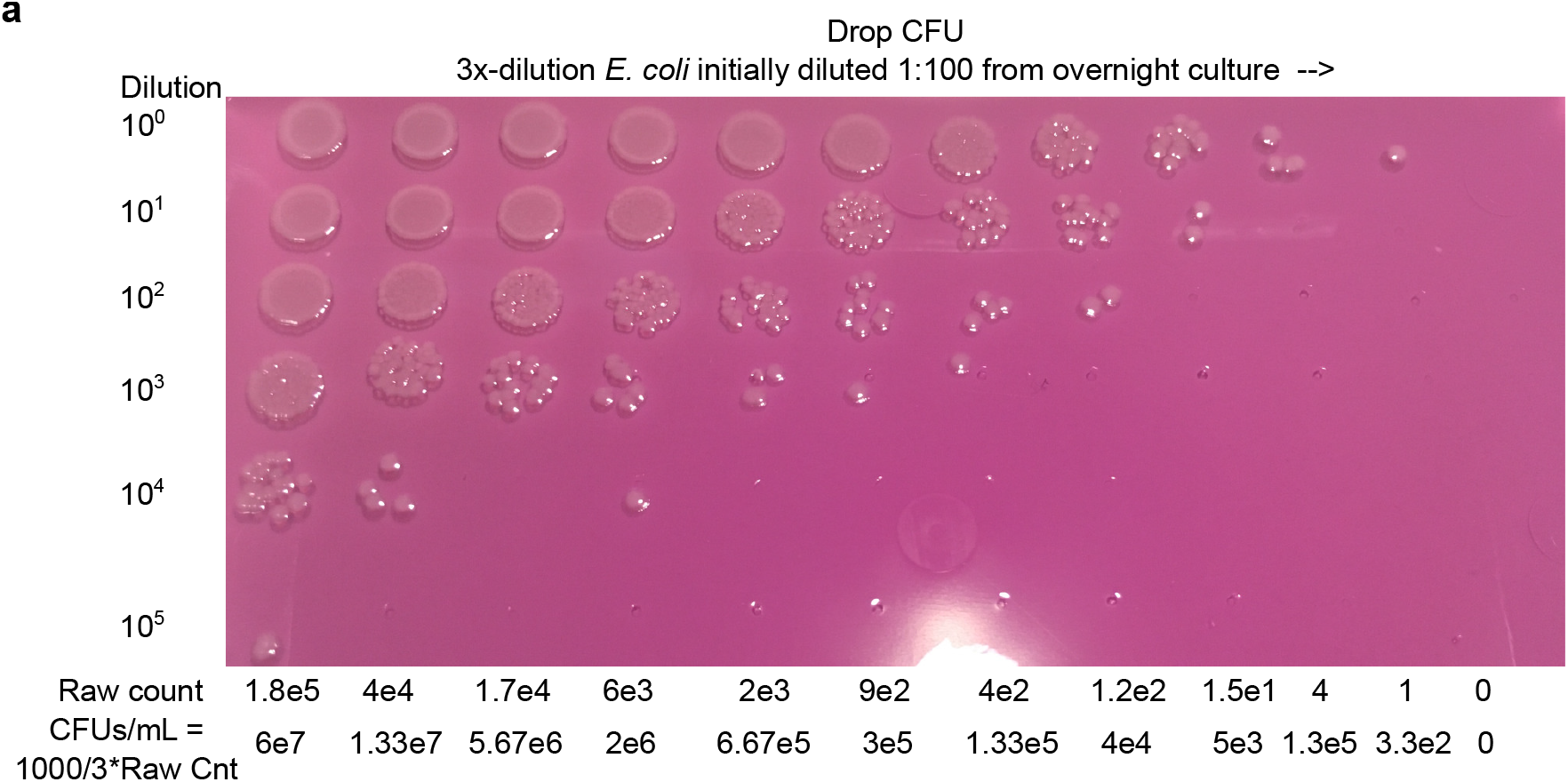
Example drop CFU plate. a) Each condition (columns) is diluted with a 10-fold serial dilution (rows) and 3uL are spotted on a 1.5% LB agar pad poured into an empty tip box. Colonies are counted for the dilution row where individual colonies are discrete. These counts are used to calculate the CFUs/mL (bottom).

**Figure S4:**
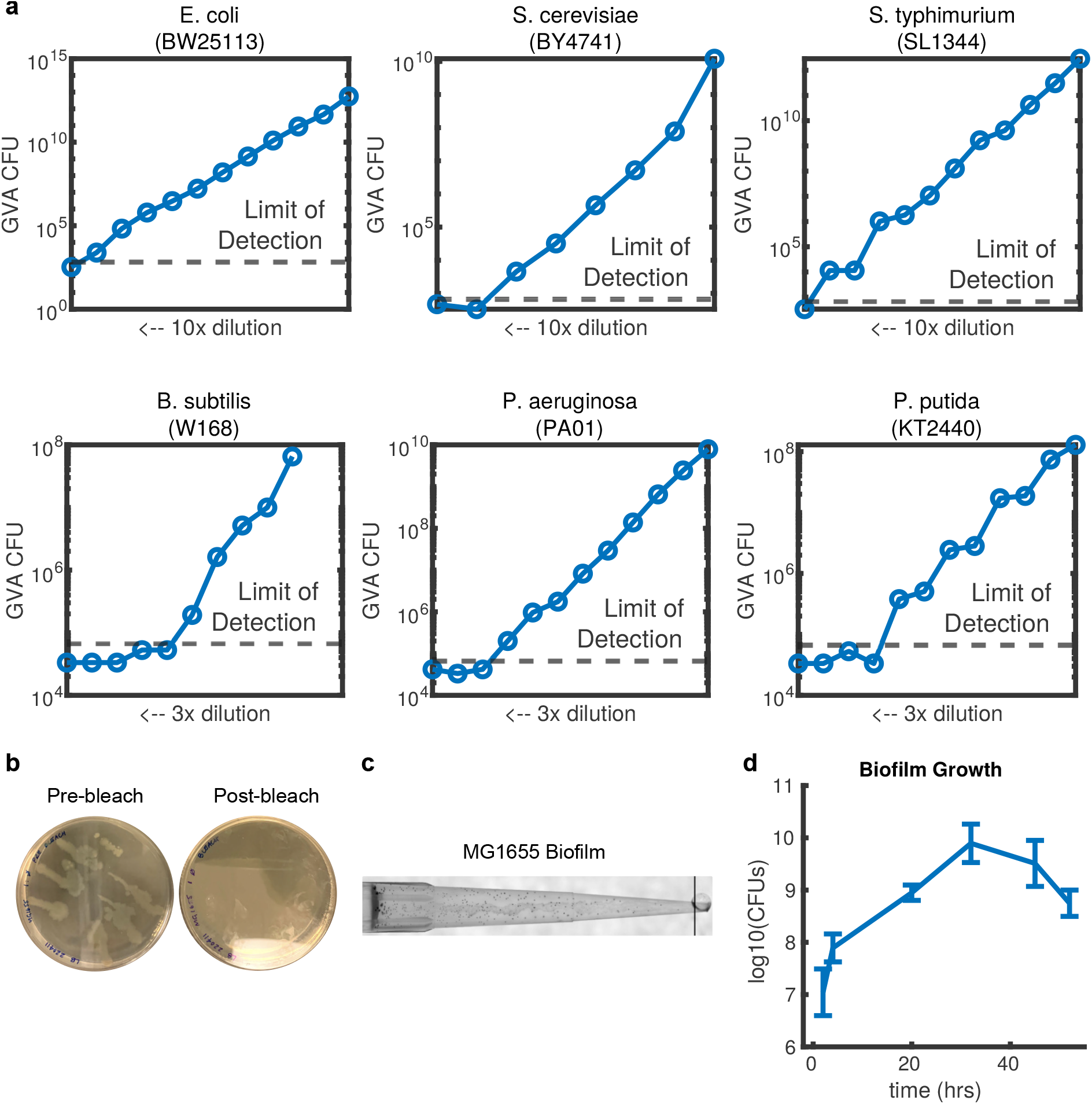
GVA calculations for different species. a) For the six species tested with GVA, the estimated number of CFUs/mL for different dilution series. b) Plates streaked with pipette tip after GVA embedding before or after bleach wash. No change in CFUs/mL were observed after bleach wash. c) Example GVA pipette tip for an *E. coli* biofilm. See Methods for culture and dissociation protocol. d) Biofilm growth over time. Errorbars correspond to standard deviation between ≥ 5 biological replicates.

**Figure S5:**
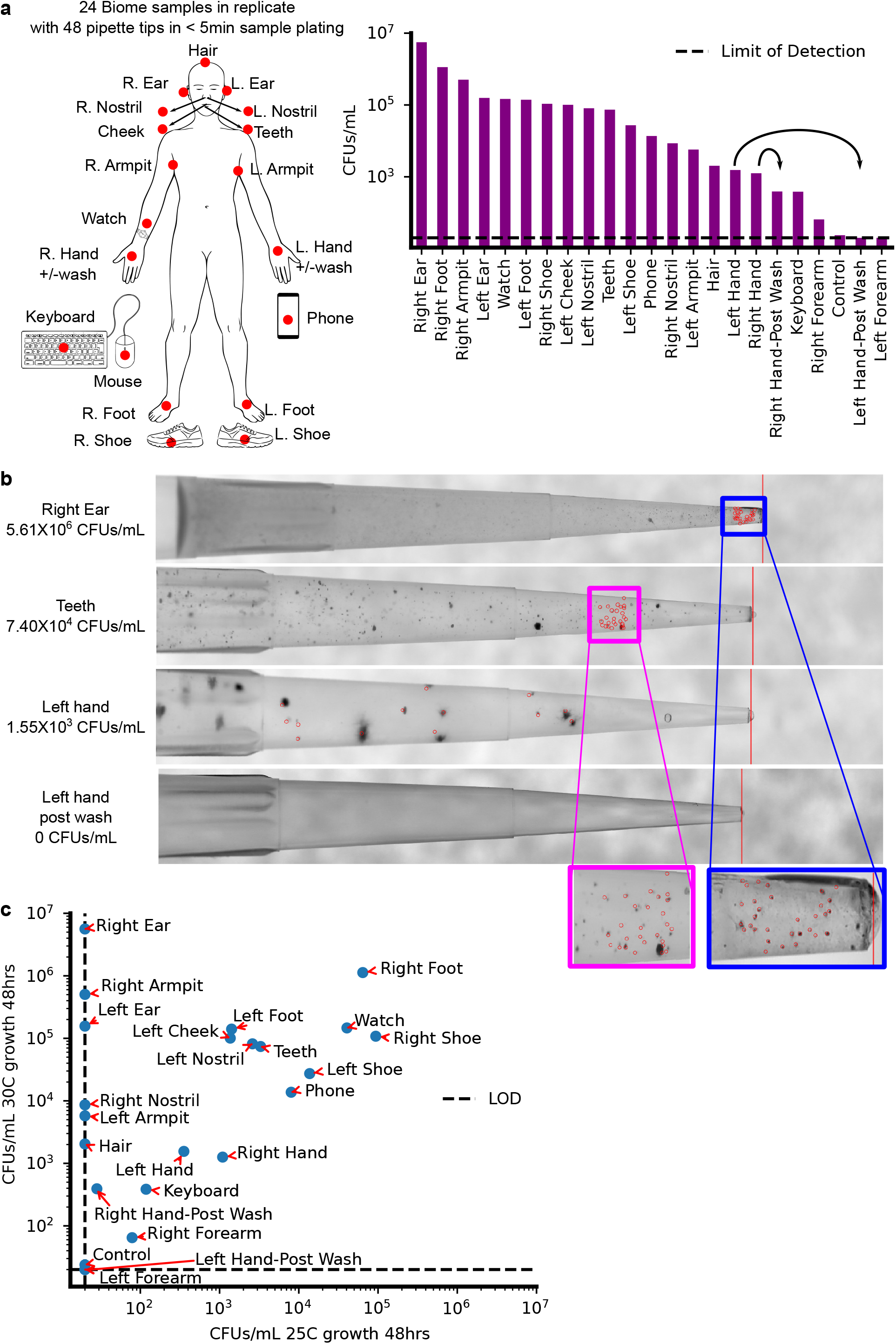
Biome sampling using GVA. a) Twenty-four positions (red dots) on a volunteer were swabbed vigorously for 15 seconds before being placed in 1 mL of LB medium and vortexed for 10 seconds. 50 *µ*L of the sample was then mixed with 150 *µ*L of 0.66% melted LB agar to a final concentration of 0.5% agar and allowed to gel in the tips. With this protocol, the lower limit of detection was 20 CFUs/mL (dotted line). The sample replicates were incubated at 30°C or 25°C for 48 hours before imaging. b) Example pipette tips for different sample regions reveals diverse colony structure and concentration for different biome locations. All samples were stained with TTC. c) Samples from higher thermal regions (ear, armpit) grew at 30°C but did not grow at 25°C indicating the temperature selectivity of different species grown in the pipette tip.

**Figure S6:**
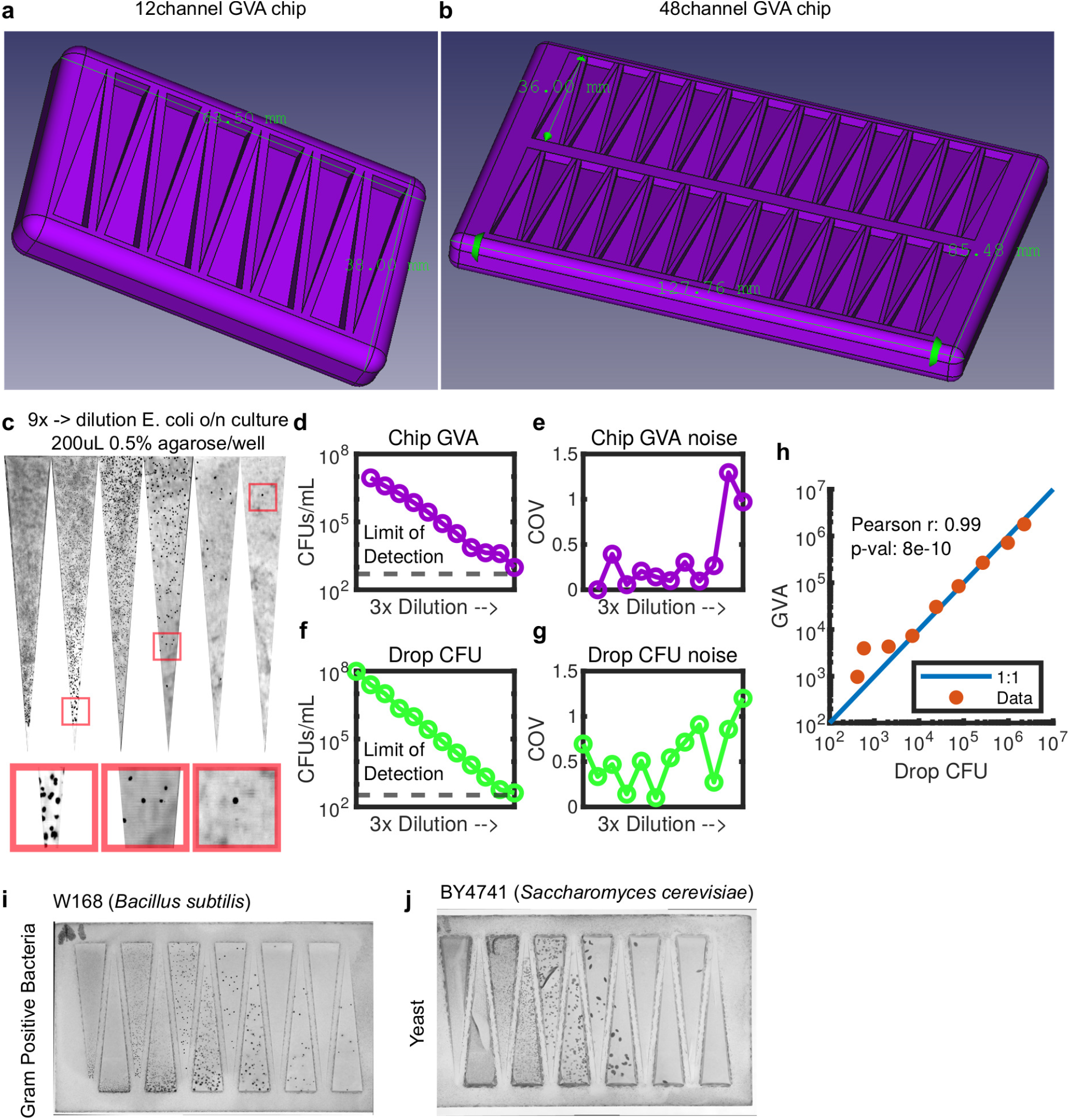
Chip version of GVA uses the square pyramid geometry. a,b) 3D printed molds for creating square pyramid for 12 (a) and 48 (b) conditions. c) Picture of a 9x dilution series of *E. coli* cultures on the GVA chip. d) GVA calculated CFUs/mL using for a dilution series. Each dot is the mean of 4 replicates. e) The noise, measured using the coefficient of variation (COV) for the chip GVA. f) Matched drop CFU quantification to conditions in (d). g) Corresponding noise analysis for drop CFU. h) Correlation between chip GVA and drop CFU over 5 orders of magnitude. i,j) Chip GVA for gram-positive (i) and eukaryotic (j) cells.

**Figure S7:**
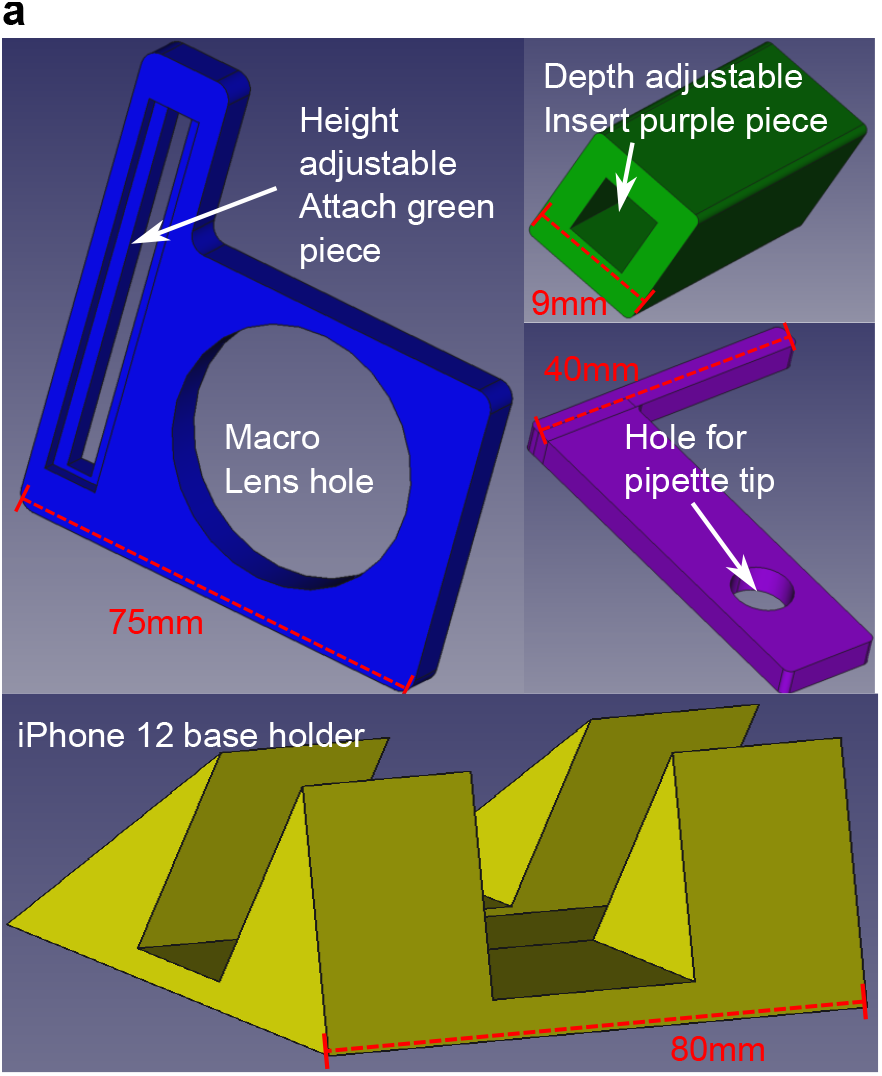
iPhone pipette tip holder. a) The 3D printed parts for stereotypically positioning a pipette tip in front of an iPhone rear camera with a Xenvo macro lens (15x magnification without the widefield lens). The blue face plate slides onto the Xenvo macro lens which is clipped to the iPhone. The green bar is attached with a screw to the side channel on the blue plate. This allows for adjusting the height by sliding the green bar in the channel. The purple extension bar slides into the green channel to adjust the imaging depth. Phone is held upright with stand (yellow). Pieces printed with standard FDM printing with PLA.

**Figure S8:**
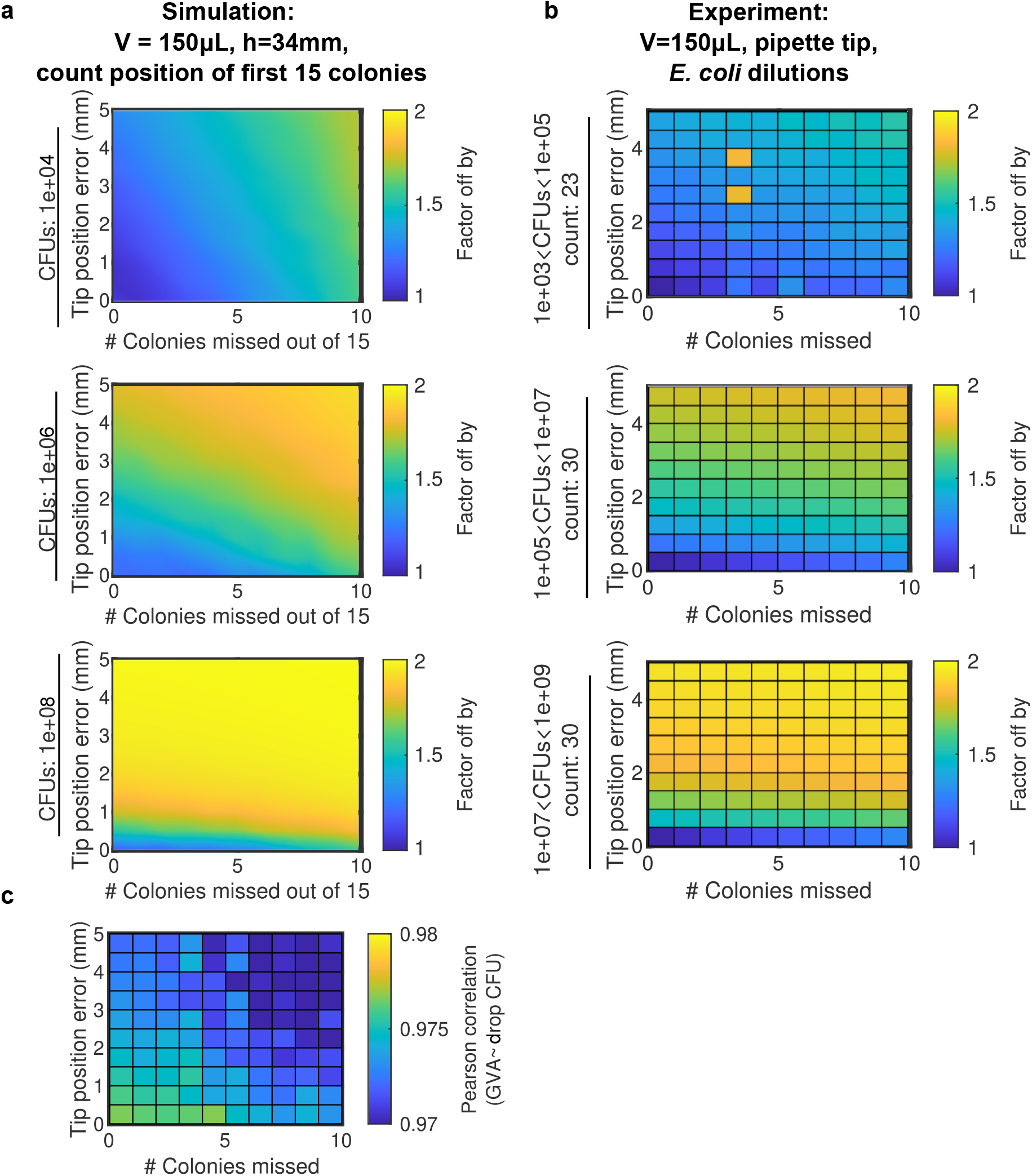
Sensitivity analysis of GVA calculations to error in missing colonies and location of the tip. a) Heatmap of the error as a function of both tip position and missing colony errors. b) Same analysis as in panel a, but with experimental data. CFUs/mL binned between 1e3 and 1e5 (top row), 1e5 and 1e7 (middle row), and 1e7 to 1e9 (bottom row). The number of pipette tips included in each bin is annotated by the count. c) Heatmap of the the Pearson correlation between the drop CFU and GVA for both tip position and missing colony errors.

**Figure S9:**
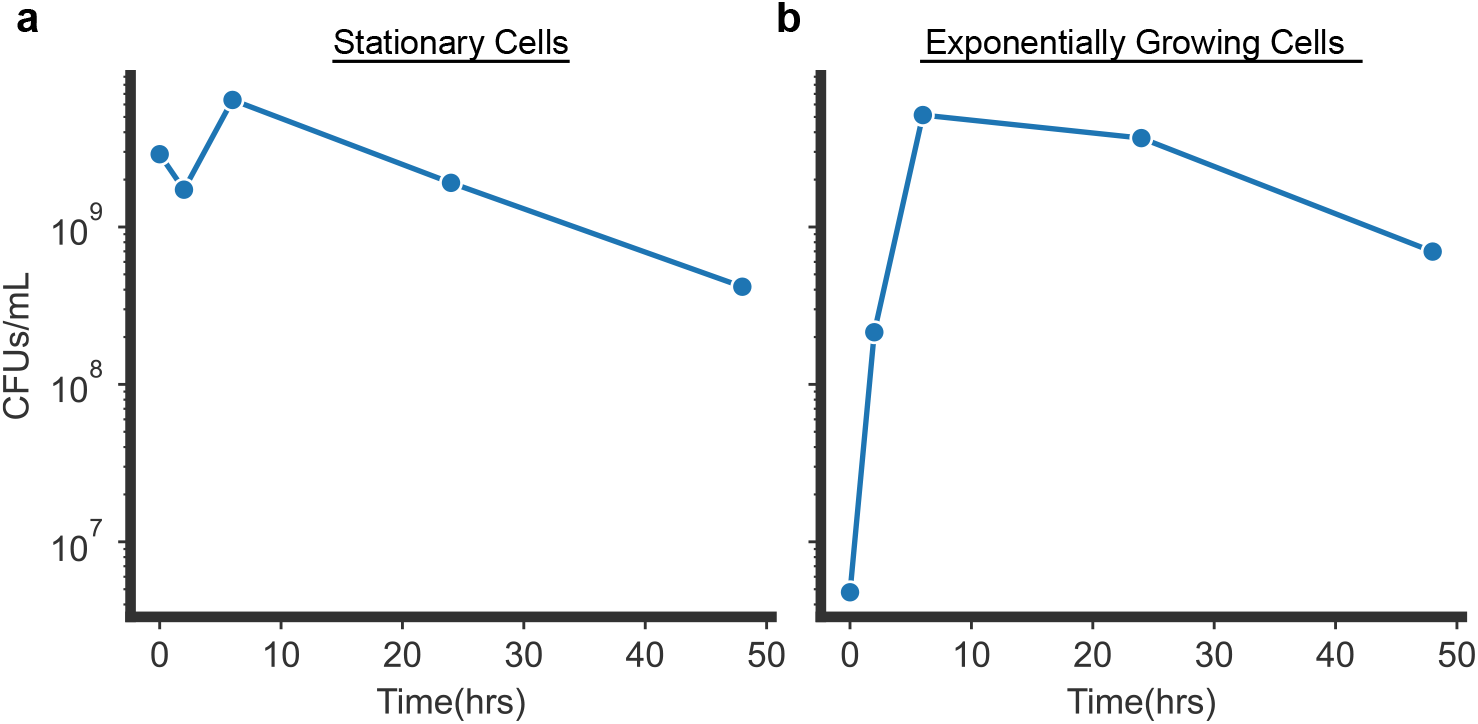
Cell counts over time in stationary versus exponential cultures. a) Number of CFUs/mL in stationary (a) versus exponential culture (b). To generate exponential culture, stationary phase cells were diluted 1:1000 in fresh LB media and place in the shaking incubator (180RPM) at 37 °C for 2 hours prior to beginning experiment.

**Figure S10:**
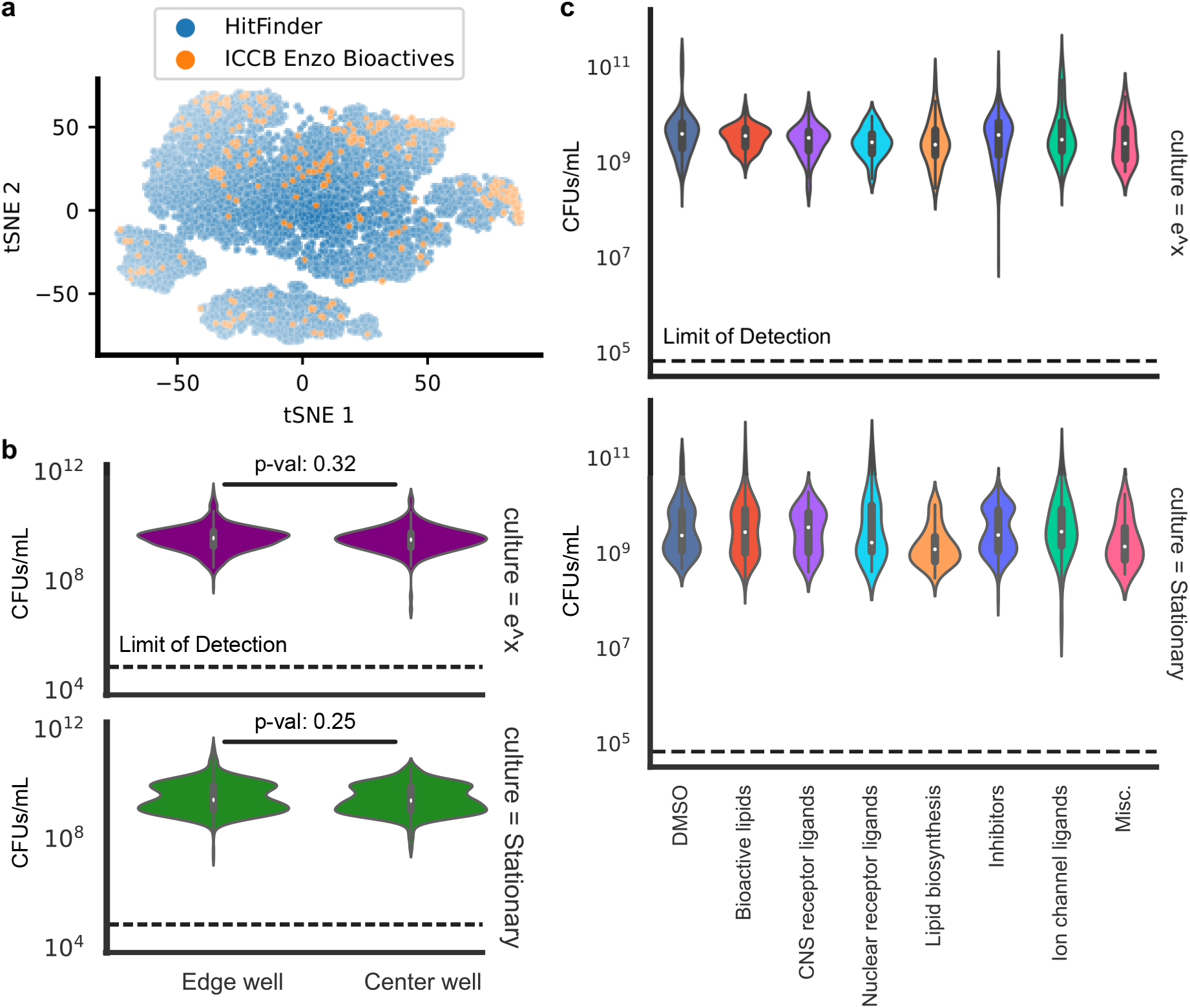
Enzo screen controls. a) Library diversity of ICCB Enzo Known Bioactive library compared to the Maybridge HitFinder library. Tanimoto similarity between all molecules based on SMILES was calculated using the RDKit package in python. From this distance matrix, the tSNE embedding was initialized with PCA and computed with a perplexity of 50. b) Distribution of CFUs/mL for conditions on the edge of the plate versus in the center wells for both stationary and exponential cultures. Statistical test used a Mann-Whitney U test for nonparametric distributions (p-val*>* 0.05). c) Distribution of CFUs/mL for different drug classes identified in the Enzo Library (See Fig. 5c). No class differences were found when using ANOVA (p-val*>*0.001, p-val corrected for multiple hypothesis testing). No differences from control were found using the Pairwise Tukey Test (p-val*>*0.01, Pairwise Tukey Test)

**Figure S11:**
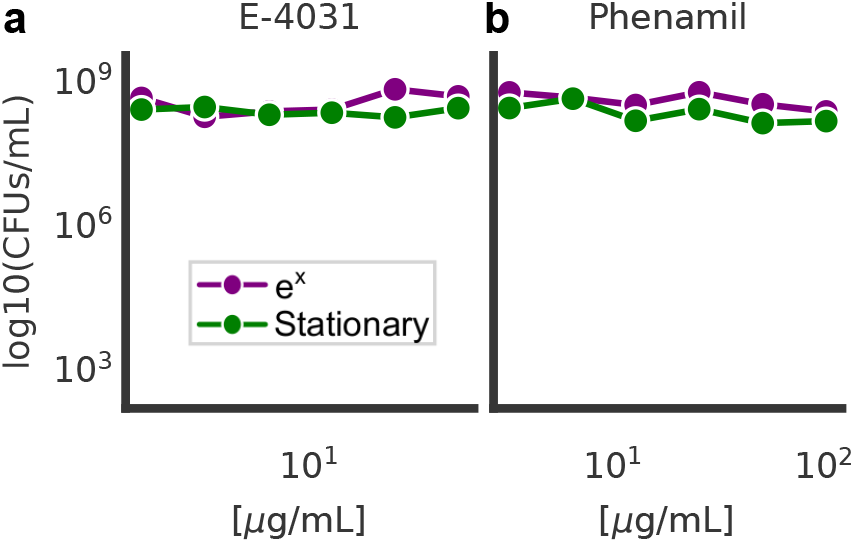
Non-validated hits from the ICCB Enzo bioactive screen. E-4031 (a) and phenamil (b) dose-response curves against stationary or exponentially (*e*^*x*^) growing cultures.

**Figure S12:**
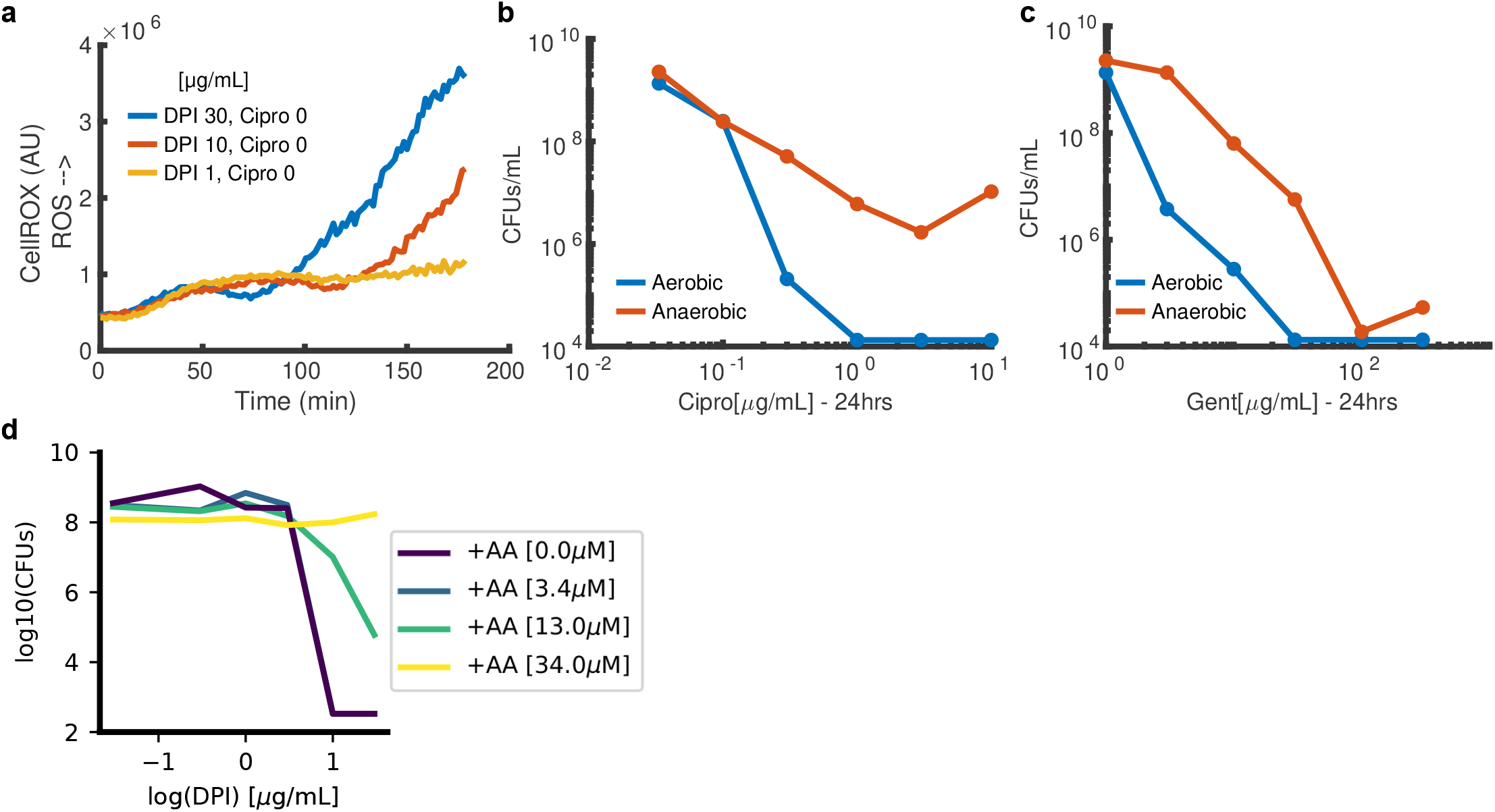
a) Duration of ROS reduction and onset of the secondary ROS spike is DPI-concentration dependent. Depicted is the median single-cell CellROX signal as a function of time for different concentrations of DPI. b,c) Dose response curve for ciprofloxacin (b) and gentamicin (c) against stationary phase cells in aerobic or anaerobic conditions. Treatment was for 24 hours. d) Efficacy of DPI as a function of increasing concentrations of the ROS-scavanger, ascorbic acid (AA).

**Figure S13:**
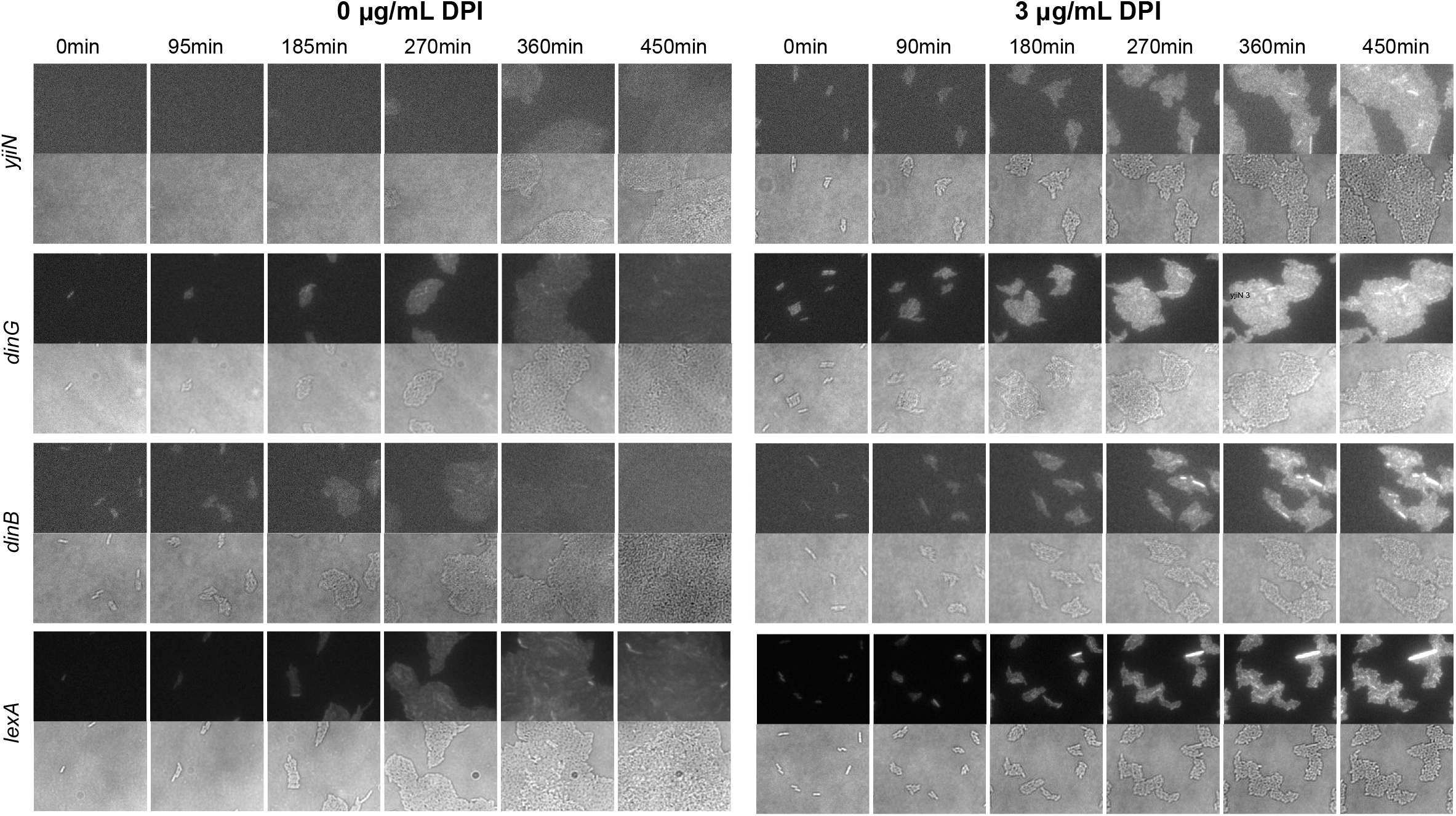
Strip charts of lexA-repressed genes (rows) using the PEC library. GFP fluorescence (top panels of each row) is proportional to each gene’s promoter activity. Bottom panel of each row depicts brightfield image. Columns correspond to different timepoints post treatment.

**Figure S14:**
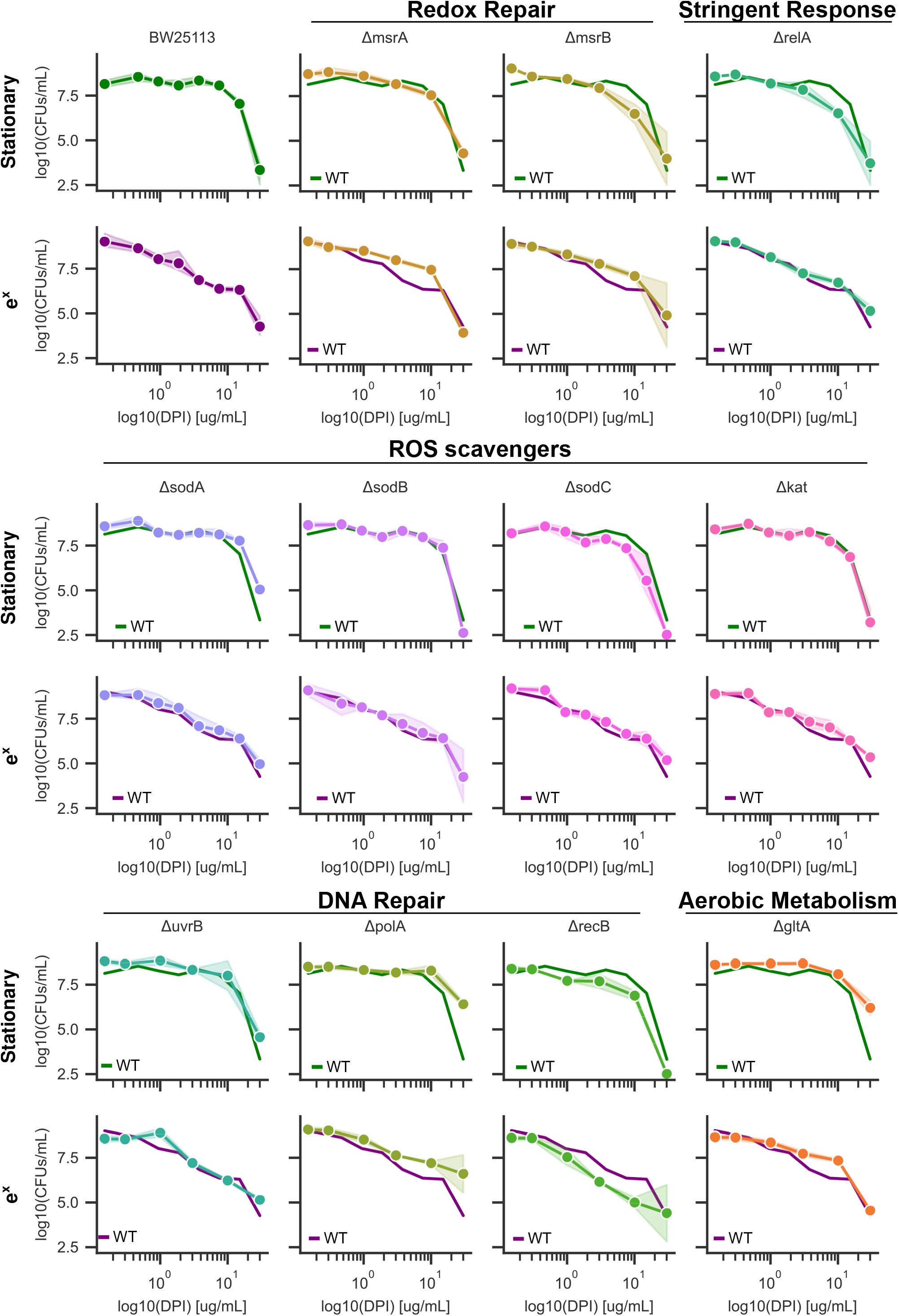
Sensitivity of gene mutants to DPI in exponential and stationary phase. Wild type reference depicted in solid line for each mutant. Error bars are the standard deviation in log space between three biological replicates. Mutants were selected from the Keio collection. Kanamycin (25 *µ*g/mL) was included in the all Keio culture conditions both in the overnight culture and during treatment with DPI to maintain gene knockout.

**Figure S15:**
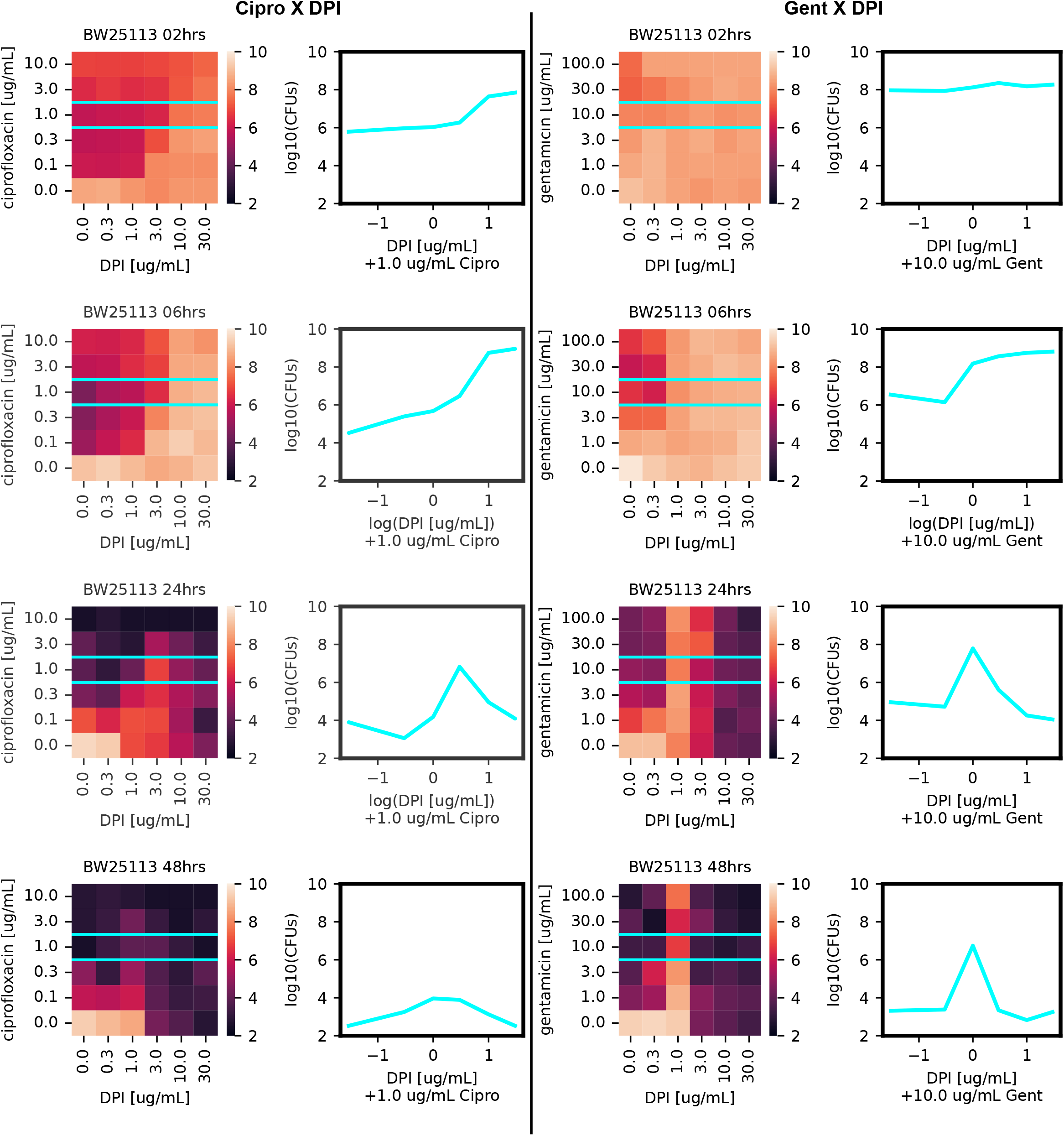
GVA temporal checkerboard of DPI crossed with either ciprofloxacin (left panels) or gentamicin (right panels) against *E. coli*. Treatment time increases down the rows. Each square in the heatmap was the mean of duplicate conditions. Colorbar correspond to the measured log10(CFUs/mL) for each combination. Left panel shows line trace (cyan) for the DPI dose response at 1*µ*g/mL ciprofloxacin or 10 *µ*g/mL gentamicin.

**Figure S16:**
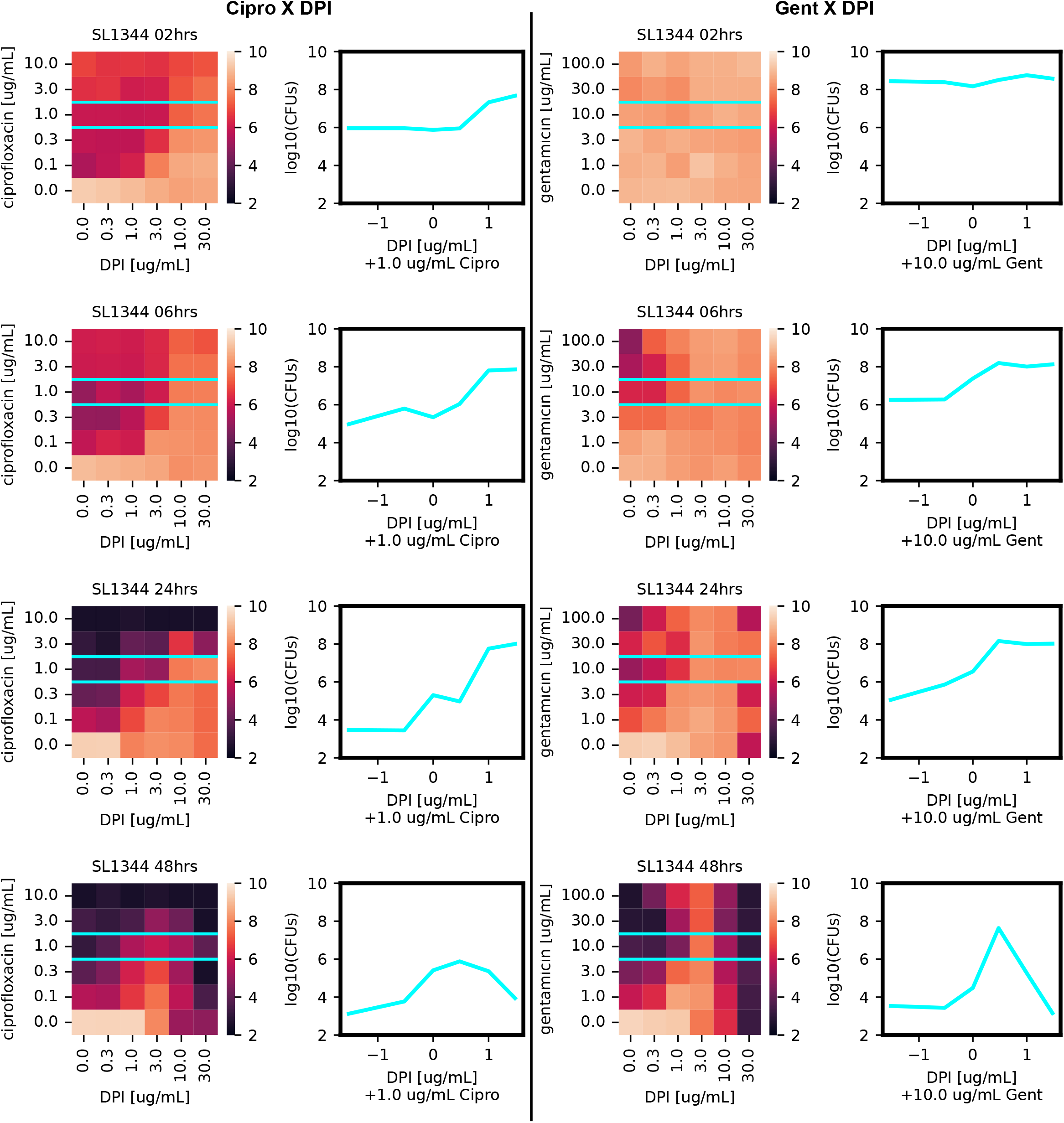
GVA temporal checkerboard of DPI crossed with either ciprofloxacin (left panels) or gentamicin (right panels) against *S. typhimurium*.

## 4 Derivation of the axial probability density function for a cone

Assuming single cells are well mixed before being suspended and cast into a 3D cone, the probability of a colony forming at distance *x* from the origin is proportional to the percent of the total volume (*V*) comprised by the infinitesimal volume (*dV*) at *x. dV* is defined as

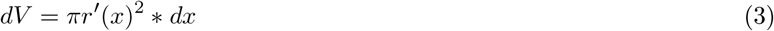

where *r*^*′*^(*x*) is the radius of the circle at *x* (Fig. S1a, cyan circle). Based on the geometry in Fig. S1a (right panel), we find

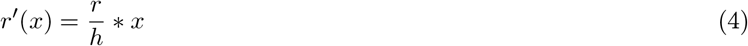

where *r* is the radius of the cone’s base and *h* is the height of the cone.

The probability density function (PDF) for this geometry can be solved for by

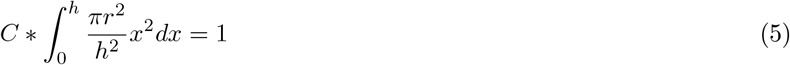

where *C* is the normalization constant and is equal to the inverse of the the volume *V* (i.e. 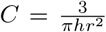) This leads to the following PDF (Fig. S1b)

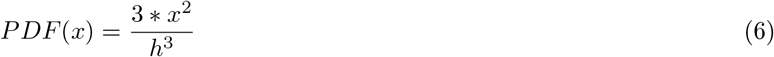

The associated Cumulative Distribution Function (CDF) can be found from the integral (Fig. S1c)

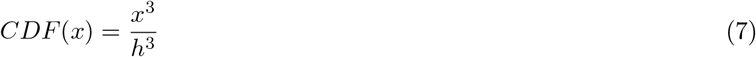

We can observe here that regardless of the base shape of the cone or pyramid, as long as it is axially symmetric, this PDF holds (Fig. S1d) as a result of the specific geometry of *dV* canceling out of the PDF due to the normalization constant. Following the same derivation, the PDF of a cylinder is found to be a constant 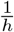 and the PDF of a 2D wedge is 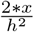 (Fig. 1b). Because of the exponent on *x*, the PDF of the cone gives the largest dynamic range in the probability (Fig. 1b). The CDF measures the likelihood of having found a colony as function of *x* if only a single colony is in the cone (Fig. 1b). The PDF is the probability of finding a colony at any point *x* for only a single colony in the cone. Therefore, there are two equivalent ways of calculating the number of CFUs using either the PDF or the CDF (Fig. S1e).

With the PDF we can estimate the number of CFUs/mL using the equation

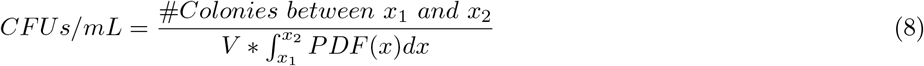

where *x*_1_ and *x*_2_ is the position of the first and last colony and *V* is the cone volume.

With the CDF,

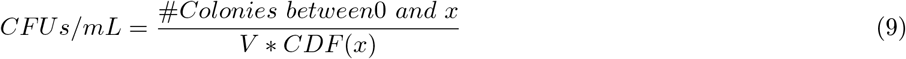

In practice, we find using the PDF estimator to be more convenient because it does not depend on identifying the first colony from the tip, but mathematically these are equivalent.

